# A Heterogeneous Graph Framework for Inference of Metabolite–Protein-Drug Interaction Networks

**DOI:** 10.64898/2025.12.18.695016

**Authors:** Yuntao Lu, Sriram Chandrasekaran

## Abstract

Metabolite–protein interactions (MPIs) are essential for coordinating cellular metabolism and signaling. Yet, MPIs remain incompletely characterized due to the limited scalability of experimental methods and the complexity of tissue-specific regulatory mechanisms. Existing computational approaches often focus on direct interactions and overlook higher-order associations and effect of drug perturbations. Here we introduce TopoMPI, a graph-based framework that integrates five types of biological relationships—metabolite–metabolite (MMI), protein–protein (PPI), metabolite–protein (MPI), drug–protein (DPI), and drug–drug (DDI)—into a heterogeneous network structure. It comprises three complementary sub-models targeting direct interaction prediction, high-order association discovery, and drug-protein-metabolite triplet interaction inference. Comprehensive evaluation across 24 tissue-specific MPI networks, protein-metabolite association studies and pharmacological metabolomic datasets confirm the biological relevance, robustness and generalizability of TopoMPI for MPI prediction with AUCs ranging from 0.79 to 0.86. TopoMPI provides a scalable framework for systems-level characterization of metabolic regulation and drug mode-of-action.

## 1. Introduction

Metabolite–protein interactions (MPIs) modulate key biological processes, including enzyme activity, signal transduction, transcriptional regulation, and intercellular communication, through covalent or non-covalent binding with proteins [3–6]. These interactions include canonical mechanisms such as metabolites acting as substrates or allosteric regulators of enzymes, as well as broader influences on non-enzymatic proteins like transcription factors, membrane receptors, and multi-protein complexes [7,8]. Consequently, systematic identification and characterization of MPIs are essential for understanding metabolic regulation across tissue contexts and for elucidating transitions between physiological and pathological states [9].

Despite the recognized importance of MPIs [10–12], their comprehensive characterization at the systems level remains limited [13]. A major challenge lies in their intrinsic complexity and dynamic nature—many MPIs are transient and occur with low binding affinity, making them difficult to capture using conventional in vitro assays [14,15]. Although recent advances in experimental technologies—such as tandem affinity purification [61], high-resolution NMR relaxometry [17], limited proteolysis-coupled mass spectrometry [18], and size-exclusion-based interaction enrichment methods [19]—have expanded the experimental toolkit for probing MPIs, these methods remain constrained by high cost, technical complexity, and limited throughput [20].

On the other hand, integrative analysis of metabolomics and transcriptomics offers higher scalability and throughput for MPI inference [21,22], but these approaches often rely on statistical co-expression patterns that may lack mechanistic interpretability [23,24].

Network-based modeling offers a new perspective for systematically predicting MPIs. These interactions are often embedded within complex biological systems [25–27], such as metabolic pathways, signaling cascades, and protein–protein interaction (PPI) networks. The presence of a specific metabolite–protein interaction is determined not only by the molecular properties of the involved entities but also by their structural roles and positions in the network—such as the breadth of reactions a metabolite participates in or the centrality of a protein in the overall topology.

Graph neural networks (GNNs), which enable end-to-end learning of node representations from graph-structured data, have demonstrated strong predictive performance across a variety of biomedical tasks [28,29], including drug–target interaction prediction, protein function annotation, and gene–disease association mapping [30–33]. Heterogeneous GNNs extend this framework by incorporating multiple types of nodes (e.g., metabolites, proteins, drugs) and diverse semantic relationships (e.g., metabolic co-participation, PPIs, drug–target links), allowing the model to capture multi-level regulatory patterns [34–37].

Recent studies have begun to apply GNNs to MPI prediction, incorporating information such as metabolite structural similarity and PPI data into heterogeneous networks for joint embedding learning [38]. Other efforts have combined protein sequence features with metabolite fingerprints through attention-based architectures to extract interaction-specific patterns [39,40], or have integrated genome-scale metabolic models to support MPI inference at a systems level [41]. These advances suggest that graph-based learning methods are highly scalable for MPI modeling.

However, three key limitations remain. First, while GNNs are structurally designed for graph inputs, many models still emphasize node-level attributes—such as protein sequences or chemical fingerprints—while underutilizing information from network paths, higher-order neighborhoods, or global topology. This restricts the model’s ability to capture structural patterns inherent to MPIs. Secondly, most existing approaches do not account for variations in interaction patterns across tissues or environmental conditions. Thirdly, MPI networks involve multiple types of entities and relations. There is no standardized framework for effectively integrating these diverse edge types, quantifying their contributions, or evaluating them through unified representation learning and interpretable assessments.

In summary, although GNNs have introduced a new modeling paradigm for MPI prediction, further development is needed to enhance their structural semantic modeling, incorporate physiological or condition-specific contexts, and support the integration of heterogeneous relationship types. To address these issues, we propose an integrated graph neural network framework, termed TopoMPI (Topological Metabolite–Protein Interaction prediction model). Through extensive evaluations, including cross-tissue generalization tests, perturbation-based robustness analysis, edge-type ablation studies, and functional consistency validation with external experimental data, TopoMPI demonstrates strong predictive performance, and the capacity to uncover biologically meaningful regulatory structures.

## 2. Results

### 2.1 Overview of Network Construction and Model Architecture

#### 2.1.1 Functional Roles of the Three TopoMPI Submodels

TopoMPI comprises three complementary submodels, namely TopoMPI-D, TopoMPI-I, and TopoMPI-C, and each submodel is designed to address a distinct aspect of metabolite–protein interaction (MPI) inference. All three models operate on a unified heterogeneous graph that integrates metabolites, proteins, and drugs as node types and incorporates multiple biological relations (**Figure 1**). Node representations are initialized using large-scale pretrained embeddings: protein sequences are encoded with ESM2 [42], and metabolite and drug structures are embedded via ChemBERTa-encoded SMILES strings [43], ensuring that all node types reside in a shared semantic space suitable for cross-modal learning.

**Figure 1.**
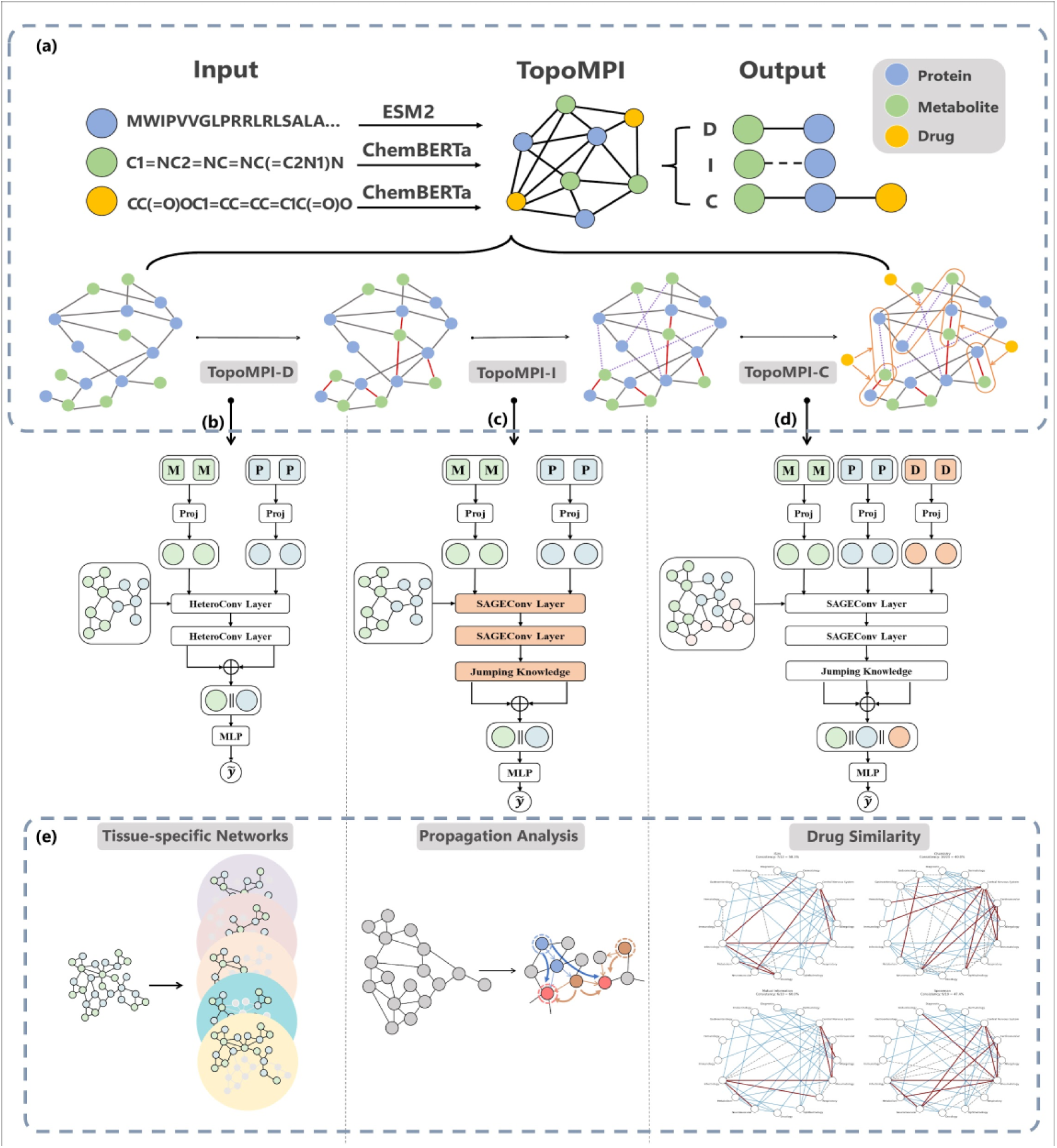
Overview of the TopoMPI framework and the functionality of its three constituent models. This figure illustrates the overall architecture of TopoMPI and the roles of its three specialized submodels in predicting metabolite–protein interactions (MPIs) under distinct biological contexts. The top panel **(a)** depicts a progression of four network types from left to right: the original background network (input graph), direct interaction prediction via TopoMPI-D, high-order structural association modeling via TopoMPI-I, and drug-dependent interaction inference via TopoMPI-C. This sequence reflects the conceptual expansion of the prediction task from direct linkage detection to context-aware, system-level modeling. The bottom portion is divided into three vertical sections, each corresponding to one model. The left column **(b)** illustrates the architecture of TopoMPI-D, which employs a Heterogeneous Graph Attention Network (Hetero-GAT) to integrate multiple edge types—including metabolite–metabolite (MMI), protein–protein (PPI), and metabolite–protein (MPI) interactions—and learn direct MPIs through localized topological features; its structure includes feature projection, dual-layer HeteroConv aggregation, and an edge-level MLP for prediction. The middle column **(c)** shows TopoMPI-I, which extends TopoMPI-D by replacing the convolution layers with SAGEConv and incorporating Jumping Knowledge aggregation to capture high -order, indirect associations between metabolites and proteins. The right column **(d)** illustrates TopoMPI-C, which further expands the model by introducing drug nodes and incorporating drug–protein (DPI) and drug–drug (DDI) interactions, thereby constructing a tri-modal heterogeneous graph to model triplet-level, drug-induced interaction paths; its architecture includes projection, convolution, and cross-modal fusion layers followed by a triplet-based MLP predictor. **(e)** Applications of the three models (see Figure 3-5). This panel summarizes the application and validation of the three models: TopoMPI-D (left) was applied to 24 tissue-specific networks for evaluating topological reconstruction (see Results 2.2); TopoMPI-I (middle) predictions were validated via comparison with protein metabolite association studies and network propagation analysis (see Results 2.3); and TopoMPI-C (right) generated drug–protein–metabolite interaction networks and enabled drug similarity-based functional annotation (see Results 2.4).

The TopoMPI framework is motivated by three assumptions: (i) molecular representations derived from protein sequences and metabolite SMILES strings capture essential biochemical properties for inferring MPIs; (ii) known network topology, including direct MPIs and auxiliary edges such as PPIs, MMIs, and drug–target associations, encodes meaningful local and global structural patterns that can guide generalization; and (iii) in the presence of drug perturbations, incorporating drug structure and drug–target links enables the model to capture context-specific rewiring of metabolite–protein associations.

TopoMPI-D (Direct Interaction Prediction) is designed to recover missing MPI edges in a static heterogeneous graph. The underlying heterogeneous molecular interaction network includes 2,375 metabolites and 6,027 proteins, connected through MMI edges from Recon3D biochemical reaction model [44], PPI edges from STRING database [45], and MPI edges from STITCH database [46]. TopoMPI-D applies relation-aware attention mechanisms to aggregate local topological information and predict edge-level interaction likelihoods.

TopoMPI-I (Indirect Association Prediction): TopoMPI-I includes the same MMI, MPI, PPI networks based on RECON3D, STRING and STITCH as TopoMPI-D, but it extends beyond direct edges by identifying higher-order or topologically unconnected MPI associations. TopoMPI-I is trained on a metabolite–protein association dataset comprising 1,365 metabolites and 974 proteins [47], deriving association scores from embedding similarity rather than edge-level classification. This enables detection of indirect or latent coupling patterns that may not be reflected in explicit network connectivity. Compared to TopoMPI-D, it replaces GATConv with SAGEConv to aggregate multiscale neighborhood information, and introduces a Jumping Knowledge (JK) mechanism to capture hierarchical structural features.

TopoMPI-C (Drug-Conditioned Triplet Prediction): TopoMPI-C introduces drug perturbation as a contextual dimension by augmenting the heterogeneous graph with DPI and DDI edges from DrugBank[48]. Building upon the same MPI datasets used for TopoMPI-I, TopoMPI-C further incorporates 890 drugs to model pharmacologically induced metabolic rewiring. Supervision from drug-perturbed metabolomics enables the model to assess whether a drug–protein–metabolite triplet constitutes a valid interaction unit under pharmacological intervention, thereby capturing condition-dependent shifts in metabolic regulation.

Collectively, the three submodels offer complementary perspectives on MPI inference: TopoMPI-D captures missing direct edges, TopoMPI-I uncovers indirect functional associations, and TopoMPI-C elucidates drug-modulated interaction pathways.

#### 2.1.2 Model Performance Evaluation

To systematically evaluate the performance of the TopoMPI models in predicting metabolite–protein interactions (MPIs), we adopted a unified testing framework. Positive samples for the test set were drawn from the STITCH database, consisting of MPIs with a combined score between 500 and 900—representing high-confidence interactions that were not used during training. For TopoMPI-D, negative samples were generated by randomly sampling metabolite–protein pairs absent from the network. For TopoMPI-I and TopoMPI-C, negative samples were directly taken from the original datasets used to train the models [47], ensuring evaluation fairness across models.

TopoMPI-D achieved the highest scores across all evaluation metrics (AUC, F1-score, Precision, and Recall), indicating its superior performance in classifying direct MPI pairs (**Figure 2a**). TopoMPI-I and TopoMPI-C exhibited slightly lower scores on certain metrics but demonstrated consistent balance between Precision and Recall, suggesting stable generalization and effective use of structural information. This performance gap is partly explained by training data scale: TopoMPI-D was trained on nearly ten times more samples than the other models. Moreover, while TopoMPI-I and TopoMPI-C share the same base model, TopoMPI-C also incorporates drug conditions, effectively increasing its training set size and yielding better performance.

**Figure 2.**
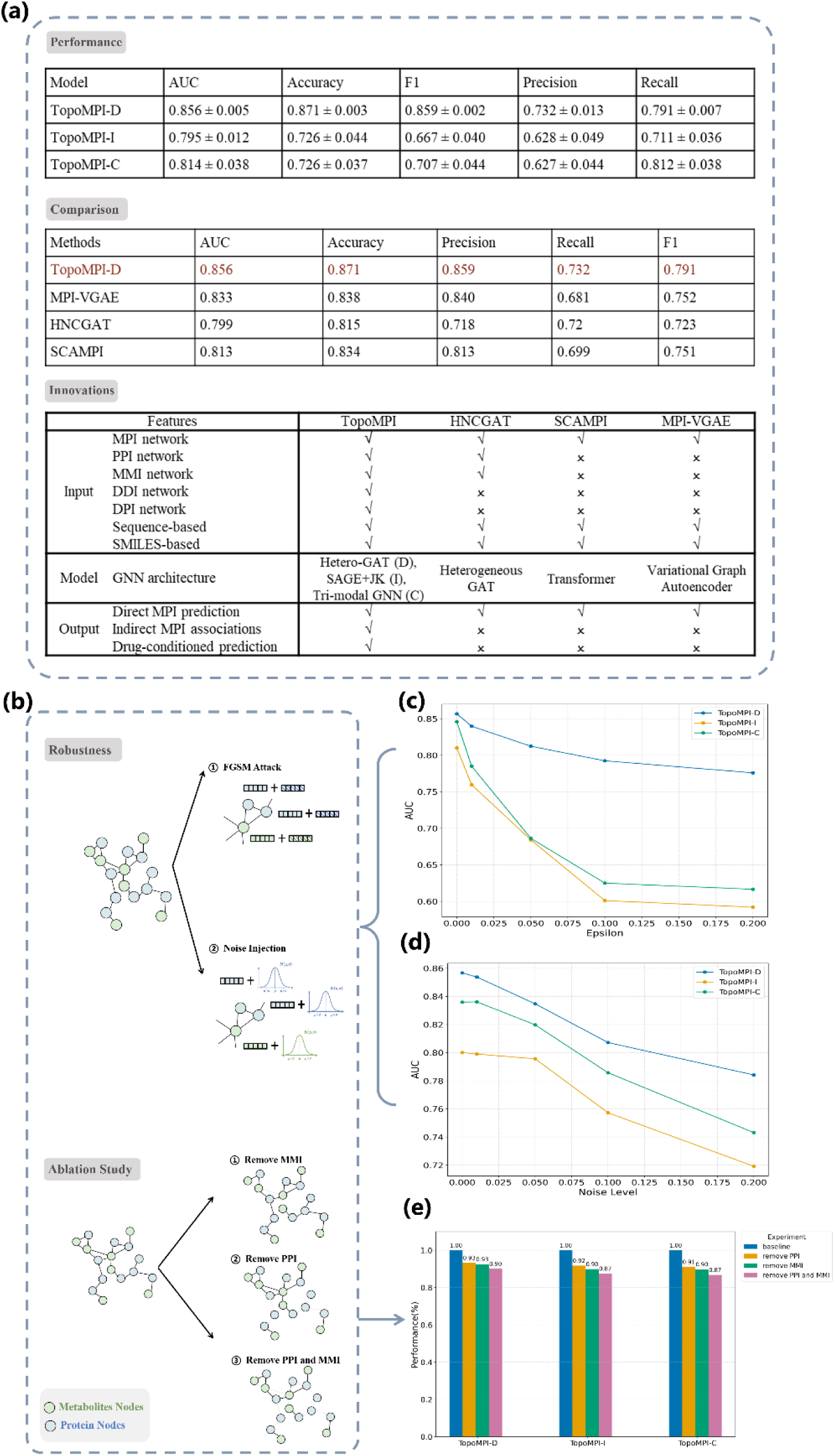
**Performance summary, robustness assessment, and ablation analysis of the TopoMPI models. (a) Summary of model performance and distinguishing features of the TopoMPI framework.**Three tables report: (i) predictive performance of the TopoMPI submodels, (ii) comparison of TopoMPI-D with representative benchmark methods, and (iii) key architectural features that differentiate TopoMPI from existing MPI prediction strategies. **(b) Schematic overview of the robustness and ablation experiments.**The robustness analysis includes Gaussian noise injection and Fast Gradient Sign Method (FGSM) adversarial perturbations, which test the stability of submodel predictions under feature-level disturbances (corresponding results shown in panels c and d).The structural ablation study systematically removes PPI, MMI, or both edge types to assess model dependency on auxiliary graph components (corresponding results shown in panel e). **(c) Robustness under Gaussian noise injection**. The x-axis indicates the standard deviation of the injected noise (Noise Level), simulating experimental or measurement-related perturbations; the y-axis shows the model’s AUC, reflecting predictive performance under different noise levels. **(d) Robustness under adversarial perturbation via the Fast Gradient Sign Method (FGSM)**. The x-axis represents the perturbation magnitude (Epsilon), and the y-axis reports the corresponding F1 score. In both tests, TopoMPI-D demonstrates the most stable performance with minimal degradation, indicating strong robustness. TopoMPI-I and TopoMPI-C show moderate performance declines under increasing perturbations but remain functionally stable, confirming the models’ general resistance to feature noise. **(e) Model performance under structural ablation experiments.** The x-axis represents different ablation groups, the y-axis represents the relative change in AUC (%) compared to each model’s baseline performance. Bar labels indicate the exact magnitude of change. All models exhibit sensitivity to graph structure integrity, with the most pronounced performance drop occurring when both PPI and MMI edges are removed. These results highlight the critical role of MMI and PPI networks in supporting accurate MPI prediction across all three models.

Since existing methods primarily focus on modeling direct interactions, we conducted comparative evaluation only for TopoMPI-D. Three representative neural-network–based frameworks were selected as baselines. HNCGAT applies type-specific attention-based neighborhood aggregation to encode node representations across a heterogeneous graph of proteins, metabolites, and functional annotations. It introduces a contrastive learning objective tailored to heterogeneous networks to preserve topological proximity among semantically distinct node types [38]. SCAMPI adopts a protein-centric strategy that uses transformer-based self- and cross-attention mechanisms to extract interaction-specific features directly from protein sequences, without explicitly modeling metabolites or network-level context [39]. MPI-VGAE, on the other hand, constructs a large heterogeneous graph of enzyme-catalyzed reactions and employs a variational graph autoencoder to learn latent embeddings for enzymatic function prediction across multiple species [40]. While all three aim to infer MPIs, TopoMPI-D differs by integrating auxiliary biological networks (PPI and MMI) and pretrained cross-modal embeddings within a unified heterogeneous space. Consistent with these design choices, TopoMPI-D outperformed all baselines across AUC, F1-score, and accuracy (**Figures 2a**).

To assess model stability under input perturbations and structural variation, we performed both robustness and ablation analyses. Under Gaussian noise injection and adversarial perturbations, all three models showed controlled performance degradation, with TopoMPI-D exhibiting the smallest drop, indicating the strongest resilience to feature-level noise (**Figure 2b and 2c**). Structural ablations further demonstrated the importance of auxiliary biological edges: removing PPI or MMI edges impaired model performance, and removing both produced the largest decline (**Figure 2d**). Across models, eliminating MMI edges consistently caused a slightly greater loss than removing PPIs, underscoring the strong contribution of metabolite–metabolite structural similarity to MPI prediction. Together, these analyses confirm that TopoMPI is robust to input uncertainty and that both PPI and MMI networks serve as essential structural priors for accurate relational modeling.

### 2.2 Assessing the Adaptability of TopoMPI-D in Tissue-Specific Networks

#### 2.2.1 Structural Adaptability of TopoMPI-D Across Physiological Contexts

To evaluate the adaptability of TopoMPI-D in modeling metabolite–protein interactions (MPIs) across diverse physiological contexts, we applied the model to background networks derived from 24 human tissues. This analysis aimed to assess the model’s robustness under structural heterogeneity and its capacity to generalize across different biological environments. Tissue-specific modeling is critical for MPI inference, as both the expression levels and functional interactions of proteins and metabolites vary substantially across tissues. A model that maintains consistent performance across these contexts would suggest strong environmental generalization and context-awareness.

The tissue-specific networks were constructed based on the Tabula Sapiens resource [49], using protein–protein interaction (PPI) maps from 24 human tissues reported by Li et al. [50] Using these PPI relationships, we subset the global MPI network to derive tissue-specific metabolite–protein graphs, and applied TopoMPI-D to each tissue network independently (**Figure 3a**). Specifically, for each tissue, we filtered the global MPI–PPI–MMI heterogeneous network by retaining only interactions involving proteins present in the tissue-specific PPI network, thereby constructing a biologically grounded and topologically consistent subgraph for each tissue.

**Figure 3.**
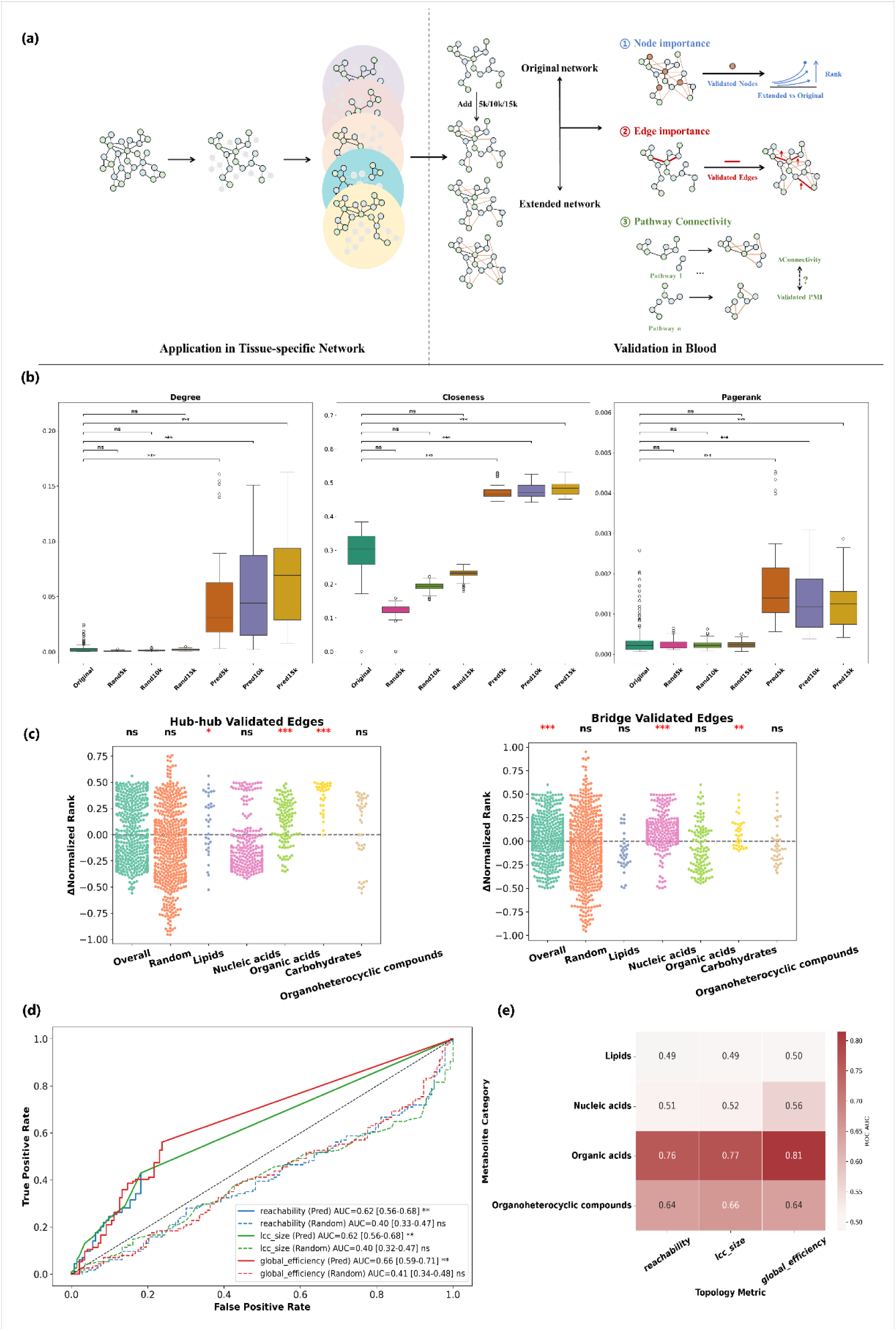
Topological adaptation and validation of TopoMPI-D predictions in tissue-specific networks. This figure presents the evaluation of TopoMPI-D’s adaptability to tissue-specific networks and the structural rationality of its predictions within the blood tissue context. subplot a illustrates the overall workflow, while subplots b–f display validation results across three analytical dimensions. **(a) Overview of the modeling and validation framework.** The upper section depicts the application of TopoMPI-D to 24 tissue-specific networks constructed using protein–protein interaction data from the Tabula Sapiens project; predictions were evaluated by comparing pre- and post-prediction networks in terms of topological changes such as node connectivity and pathway coverage. The lower section outlines the validation procedure for the blood tissue network: based on experimentally supported metabolite–protein interactions, three expanded networks (Top-5000, Top-10000, Top-15000) were generated by progressively adding high-confidence predicted edges. These networks were evaluated from three perspectives—node centrality, edge structural roles, and pathway-level connectivity. **(b) Changes in centrality of validation-set protein nodes.** This panel presents three centrality metrics—degree, closeness, and PageRank—for experimentally validated protein nodes across four network states (original and three expanded networks with increasing numbers of predicted edges). The x-axis represents network groups, and the y-axis denotes the corresponding centrality values. All three metrics exhibit consistent upward trends following edge augmentation, indicating that the model preferentially reinforces the structural prominence of biologically validated proteins. **(c) Changes in the structural roles of validation-set edges.** The plot summarizes changes in normalized rank for validated edges grouped by metabolite category. The left panel displays hub–hub edges (connecting two high-centrality nodes), and the right panel shows bridge edges (linking distinct modules). The x-axis indicates metabolite-based edge groupings, the y-axis shows the change in normalized structural importance (ΔNormalized Rank). Results suggest that predicted edges reinforce structurally significant links, with distinct patterns across metabolite classes. **(d) Discriminative power of pathway connectivity changes (ROC analysis).** Each ROC curve corresponds to a different ΔMetric—reachability, largest connected component (LCC) size, and global efficiency—used to quantify pathway-level topological improvements after prediction. The analysis evaluates whether these metrics can distinguish pathways that contain at least one validated metabolite–protein pair from those that do not. For each curve, the AUC value and its 95% bootstrap confidence interval are displayed, along with a statistical significance star (*p < 0.05; **p < 0.01; ***p < 0.001). All metrics show AUC values exceeding 0.5, with reachability and LCC size achieving significant discrimination, suggesting that TopoMPI-D contributes to biologically meaningful topological enhancements. **(e) Functional specificity of connectivity improvements across metabolite classes.** ROC-AUC values were calculated for pathway subsets grouped by metabolite class. The y-axis lists metabolite categories; the x-axis shows AUC scores for each class. Results indicate particularly high predictive performance in pathways associated with carbohydrates and organic acids (AUC > 0.7), demonstrating TopoMPI-D’s strong ability to recover structurally and functionally meaningful connections in certain metabolic contexts.

For each tissue, we generated four networks: the original graph containing only known MPI edges from databases, and three extended versions with the top 5,000, 10,000, or 15,000 highest-scoring predicted MPI edges added to the original structure. This allowed us to examine how different volumes of predicted interactions affected the network topology.

Assessment of topological features showed that tissue-specific networks supplemented with TopoMPI-D predictions displayed substantially improved connectivity (**Supplementary Figure 1**), indicating consistent structural gap-filling across tissues. These additions also enhanced biological coherence: key proteins and metabolites exhibited increased centrality in all 24 tissues, with 12.6% of hubs gaining rank across at least six tissues and most (75.7%) displaying tissue-restricted increases. The shifts aligned with known physiology, including heightened prominence of lung potassium channels and SIRT3 [51,52], CHRNA1 in muscle [53], polyphenols and glycosylated metabolites in skin [54–56], and 4-hydroxybenzoic acid in liver metabolism [57]. Full hub dynamics are reported in **Supplementary Table 1 and 2**.

We further evaluated the biological plausibility of these predictions by analyzing the connectivity of KEGG metabolic pathways before and after prediction. For each tissue, we quantified changes in pathway connectivity to assess whether the predicted MPI edges improved the structural coherence of known metabolic pathways. The results revealed tissue-specific differences in pathway connectivity gains (**Supplementary Figure 2 and Supplementary Table 3**), suggesting that the model captures distinct structural semantics in each tissue. Quantitatively, 23.8% of tissue–pathway combinations showed increased connectivity after prediction. At the tissue level, most tissues exhibited modest but consistent average gains, with 16–36% of pathways per tissue demonstrating positive improvement. A Wilcoxon signed-rank test comparing pre- and post-prediction reachability values across all pairs confirmed that these gains were highly significant (p < 1e-10). Although not evidence of direct causality, these findings indicate that TopoMPI-D systematically enhances pathway connectivity rather than generating random fluctuations, thereby supporting its ability to adapt to diverse structural backgrounds (**Supplementary Table 4**).

Importantly, several representative pathways exemplify how these gains align with tissue-specific physiology. The fat digestion and absorption pathway showed pronounced improvements in pancreas and lymph node, consistent with the pancreas as the source of lipases and cofactors essential for lipid hydrolysis, and the lymphatic system as the primary route of chylomicron transport [58–60]. Glutathione metabolism exhibited marked gains in salivary gland, bladder, and small intestine, reflecting the central role of glutathione in antioxidant defense across oral epithelia, urothelium, and intestinal mucosa [61–63]. Likewise, histidine metabolism displayed consistent improvements in lymph node, small intestine, and kidney, coherent with the conversion of histidine to histamine by immune and enterochromaffin-like cells in lymphoid and intestinal tissues, and with the kidney’s role in histidine dipeptide transport and degradation [64–66]. Together, these examples illustrate that TopoMPI-D not only improves global pathway connectivity but preferentially augments networks directly relevant to each tissue’s functional specialization, further underscoring the biological plausibility of its predictions.

#### 2.2.2 External Validation of TopoMPI-D Predictions

We leveraged a high-confidence set of experimentally validated MPIs derived from large-scale multi-omics profiling of human plasma reported by Benson et al. [67].In that work, the authors performed large-scale mass spectrometry-based metabolomics and aptamer-based proteomics on plasma samples from thousands of individuals across three independent cohorts (JHS, MESA, and HERITAGE), enabling systematic correlation analysis between circulating metabolites and proteins measured from the same biological samples. After mapping the nodes between our constructed blood network and the external dataset, we obtained 11,103 metabolite–protein association**s** that could in principle be evaluated in our network. Among these, 304 associations (2.7%) were already present in the original network, while 5,971 (53.8%) were successfully recovered by TopoMPI-D predictions. These enrichments were highly significant compared to random expectations (empirical and permutation p-values < 0.05), confirming that TopoMPI-D preferentially recovers biologically validated MPIs rather than random links.

Nodes involved in externally validated MPIs showed consistent increases in degree-, closeness-, and PageRank-based centrality after adding TopoMPI-D predictions, with PageRank exhibiting the strongest gains (**Figure 3b**). These shifts indicate that the model preferentially reinforces biologically relevant nodes rather than uniformly inflating connectivity. Validated edges also moved into more influential structural roles, either as hub–hub links or module-bridging connections, with metabolite classes displaying distinct patterns (e.g., carbohydrates gaining in both categories, lipids and organic acids mainly in hub–hub roles, nucleic acids in bridging roles; **Figure 3c**).

At the pathway level, changes in reachability, largest connected component size, and global efficiency discriminated KEGG pathways containing validated MPIs from those without (AUC up to 0.66; **Figure 3d**), with carbohydrate- and organic acid–enriched pathways showing the strongest separation (AUC > 0.7; **Figure 3e**). Together, these results demonstrate that TopoMPI-D induces targeted and biologically coherent restructuring at node, edge, and pathway scales.

Collectively, these three levels of validation — node centrality, edge topology, and pathway connectivity—consistently support the biological plausibility of TopoMPI-D’s predictions in the blood tissue context, highlighting its ability to reconstruct metabolite–protein interaction networks in a structurally and functionally meaningful manner.

### 2.3 Structural Complementarity of TopoMPI-I in Functional Protein Association Modeling

#### 2.3.1 Evaluation of TopoMPI-I via External Tissue-Specific PPI Networks

Considering that many regulatory events in metabolic systems may occur through indirect or higher-order interactions, we introduced TopoMPI-I to infer potential topological associations between metabolites and proteins that are not directly connected. To systematically evaluate whether TopoMPI-I provides meaningful structural complementarity to TopoMPI-D, we designed an indirect validation framework leveraging external protein–protein interaction (PPI) data. Specifically, we employed tissue-specific tumor-derived PPI networks curated by Laman Trip et al. [68], which provide probabilistic interaction estimates based on proteomic and transcriptomic co-regulation patterns.In that study, the authors integrated large-scale proteomics and paired transcriptomics datasets collected from 50 cancer studies, encompassing 5,726 tumor samples and 2,085 matched adjacent normal tissue samples. Protein–protein association scores were computed by quantifying transcriptional and post-transcriptional co-regulation, particularly among protein complex members.For our analysis, we selected four tissues—blood, liver, lung, and kidney—that overlapped with those used in our 24-tissue MPI framework to ensure contextual consistency. The resulting tissue-specific PPI edges from Laman Trip et al. served as independent external references to assess whether metabolite–protein pairs inferred by TopoMPI-I were topologically connected through biologically plausible protein interaction pathways (**Figure 4a**).

**Figure 4.**
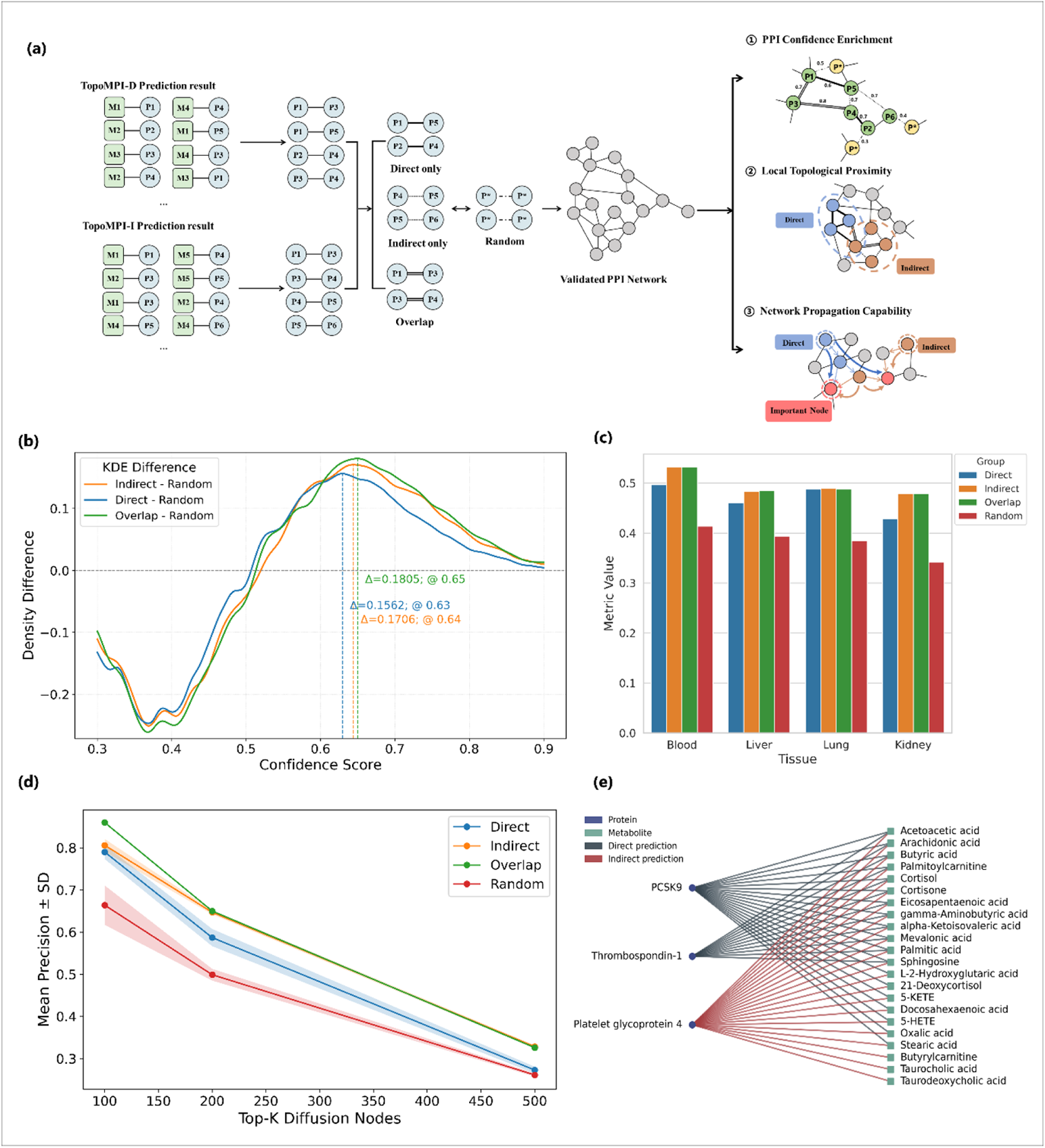
**Structural evaluation of TopoMPI-I predictions across tissue-specific networks**. This figure illustrates the evaluation of TopoMPI-I’s indirect prediction results through multi-perspective structural validation and case-based biological interpretation. Predictions are compared to those of TopoMPI-D and validated against tissue-specific PPI networks from four independent sources (blood, liver, lung, kidney). Validation is organized into three structural analyses, followed by case studies in the blood network. **(a) Overview of validation design.** The left subplot shows the construction of four prediction groups by mapping TopoMPI-D and TopoMPI-I outputs onto external PPI networks: Direct, Indirect, Overlap (Direct & Indirect), and a randomly sampled control group. The right subplot outlines three validation perspectives: (1) PPI Confidence Enrichment evaluates the distribution of confidence scores for protein pairs within each group; (2) Local Topological Proximity assesses clustering and modularity of predicted nodes; (3) Network Propagation Capability measures the ability of each group’s proteins to reach core functional hubs through random walk simulations. **(b) PPI confidence enrichment in blood.** Kernel density estimates (KDEs) show the distribution of external confidence scores for protein pairs across four groups. Indirect and Overlap groups exhibit right-shifted density peaks relative to Direct and Random, indicating that TopoMPI-I preferentially recovers high-confidence interactions overlooked by direct predictions. **(c) Topological proximity via transitivity.** Subgraphs formed by each prediction group in the blood network are compared using transitivity as a metric of local clustering. All predicted groups exceed the random baseline, with Indirect achieving the highest clustering tendency, suggesting greater structural coherence. **(d) Network propagation analysis in blood.** Using Random Walk with Restart (RWR) on a high-confidence PPI network, Precision (K = 100, 200, 500) is computed based on overlap with top PageRank nodes. Indirect predictions consistently achieve higher Precision (K = 100 and 200) than Direct-only, indicating stronger embedding in core regulatory regions. **(e) Case studies in the blood tissue network: lipid metabolism.** An example of an Indirect-only interaction in the blood network highlights a biologically validated but structurally non-obvious protein–metabolite pair. This demonstrates TopoMPI-I’s ability to recover meaningful interactions that are inaccessible to direct topology-based models.

The validation was grounded in the following core assumptions: (1) if TopoMPI-I predicts biologically relevant MPIs, the connected proteins should show higher likelihood of interaction within an independent PPI network; (2) these proteins should exhibit cohesive topological organization; and (3) signals originating from these proteins should propagate efficiently to biologically central nodes, indicating strong network diffusion potential.

Across all tissues, protein pairs derived from TopoMPI-I showed stronger enrichment for high-confidence PPIs than those derived from TopoMPI-D, with both the Indirect-only and Overlap groups displaying clear right-shifted score distributions relative to degree-matched controls (**Figure 4b; Supplementary Figure 4**). These enrichment patterns indicate that TopoMPI-I preferentially identifies metabolite-linked protein relationships with greater biological support, extending the interaction space beyond direct-edge topology.

Structurally, Indirect-only and Overlap predictions formed more cohesive subnetworks, showing higher modularity and clustering than Direct-only predictions and more efficient propagation toward central regulatory hubs, particularly in blood and kidney (**Figure 4c and 4d; Supplementary Figure 5****and 6**). This superior convergence onto biologically central nodes demonstrates that TopoMPI-I captures functionally meaningful, topologically indirect associations that remain inaccessible to direct interaction modeling.

#### 2.3.2 Complementary Insights from TopMPI-I in the Blood Network

In the blood network, TopoMPI-I provided substantial complementary biological insight beyond direct-edge prediction. Using the plasma dataset from Benson et al. study described earlier [67], TopoMPI-I recovered 2,187 validated metabolite–protein associations (19.7%) missed by TopoMPI-D, reflecting the strength of its indirect inference mechanism.

This structural advantage was evident across multiple metabolic contexts in **Figure 4e and Supplementary Figure 7**. In lipid metabolism, TopoMPI-D identified core interactions involving PCSK9 and THBS1 [69,70], whereas the multiscale neighborhood aggregation in TopoMPI-I enabled the discovery of additional CD36-centered interactions supported by experimental evidence [71]. In nucleic acid metabolism, TopoMPI-I resolved finer regulatory distinctions within the NME protein family by capturing NME1-associated metabolites that complement NME2 and ENTPD1 interactions detected by TopoMPI-D [72]. For blood regulators such as APOE and CD36 [73–75],

TopoMPI-I further revealed numerous validated metabolite associations, including more than twenty linked to CD36 that were not retrieved by direct inference [76–78]. These findings demonstrate that the capacity of TopoMPI-I to integrate information across indirect paths and higher-order neighborhoods translates directly into improved recovery of biologically grounded MPIs and a more comprehensive depiction of plasma metabolic regulation.

### 2.4 System-Level Mapping of Drug–Protein–Metabolite Interactions with TopoMPI-C

#### 2.4.1 Tripartite Drug–Protein–Metabolite Network Inference

Drug treatments frequently induce systemic responses via protein targets, leading to widespread changes in metabolite levels. To extend MPI modeling into perturbed biological contexts, we proposed TopoMPI-C—a heterogeneous graph-based framework that integrates drugs, proteins, and metabolites to capture tripartite associations under drug intervention.

Based on TopoMPI-C predictions, we constructed a three-layer interaction landscape encompassing drug categories, protein functional pathways, and metabolite classes, visualized using a Sankey diagram (**Figure 5a and 5b**). This diagram delineates the systematic structures by which distinct drug classes influence downstream metabolic modules through specific protein pathways.

**Figure 5.**
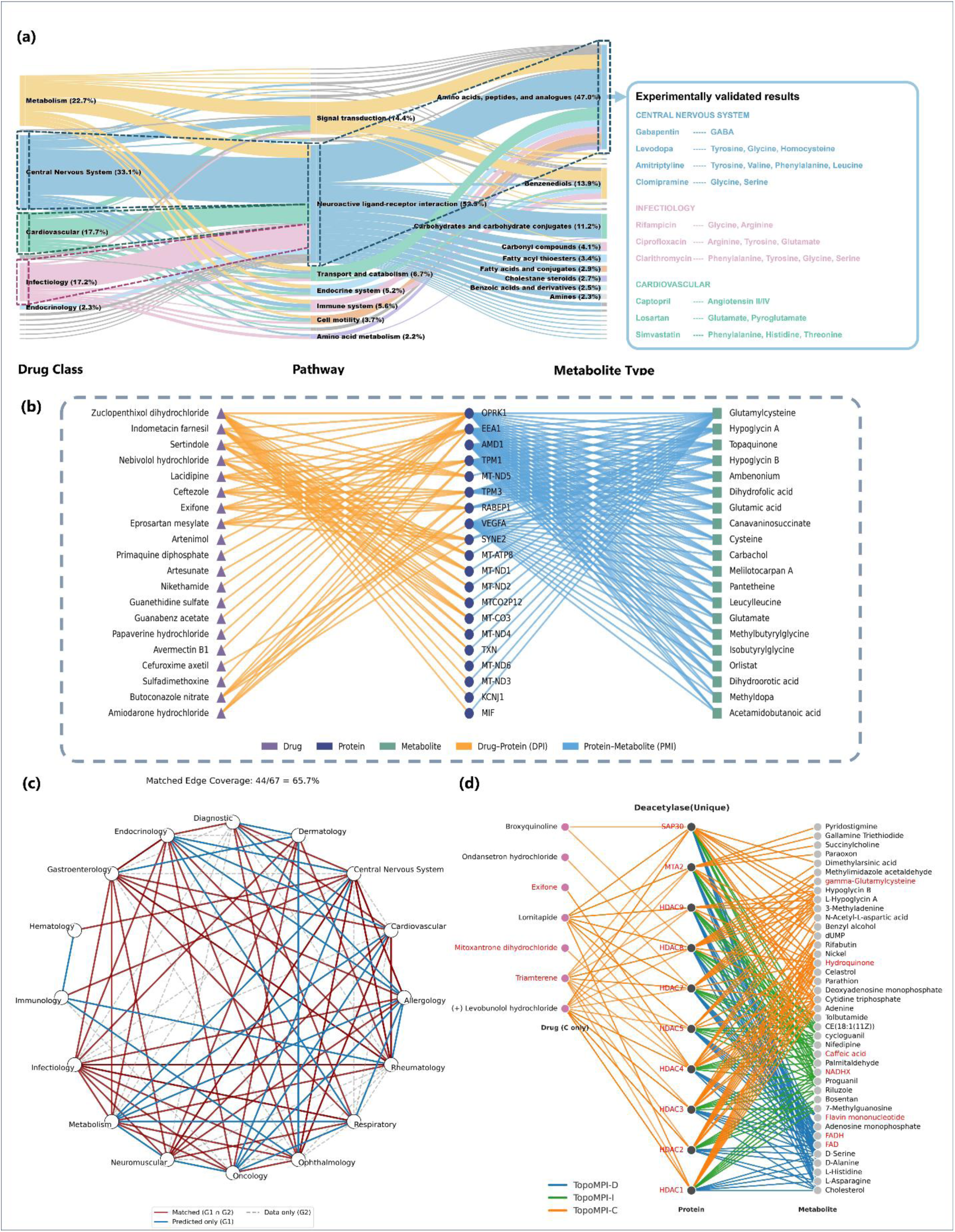
Evaluation of TopoMPI-C predictions. This figure illustrates TopoMPI-C’s capability to model drug–protein–metabolite regulatory relationships and validates the inferred drug category similarity using external benchmarks. Subplots a–d show the structural characteristics and similarity patterns learned by the model, while subplot e evaluates the consistency between predicted and established drug relationships. **(a) Sankey diagram of the three-layer regulatory landscape.** The diagram visualizes the hierarchical mapping from drug categories to protein pathways to metabolite classes. It reveals drug-specific patterns of pathway convergence and metabolite pathway reuse, reflecting multi-level regulatory mechanisms captured by TopoMPI-C. **(b) Examples of predicted tripartite interactions.** This panel provides an illustrative case of drug–protein–metabolite regulatory links inferred by TopoMPI-C, complementing the Sankey diagram by showing a concrete instance of the model’s tripartite predictions. **(c) Consensus similarity network of drug categories.** The network compares TopoMPI-C–predicted similarities with established pharmacological similarity metrics, including iSIM (integrated similarity), chemical structure similarity, mutual information, and spearman correlation. Nodes represent drug categories; edges indicate predicted similarity, with edge colors denoting consistency with external benchmarks. Blue edges highlight novel similarity relationships predicted solely by TopoMPI-C. The model recovers approximately 65.7% of known drug category similarities and reveals new, biologically plausible regulatory relationships not captured by existing methods, demonstrating TopoMPI-C’s potential to uncover latent functional connections. **(d) Unique TopoMPI predictions for histone deacetylases.** This panel shows metabolite, protein, and drug associations uniquely predicted by each TopoMPI submodel, with TopoMPI-D, -I, and -C represented by blue, green, and orange edges, respectively. Epigenetic metabolites, histone proteins, and relevant drugs are highlighted in red. These non - overlapping predictions reveal distinct regulatory inputs to deacetylase activity, spanning direct metabolic interactions, indirect redox- and signaling-linked associations, and drug-conditioned effects.

**Figure 6.**
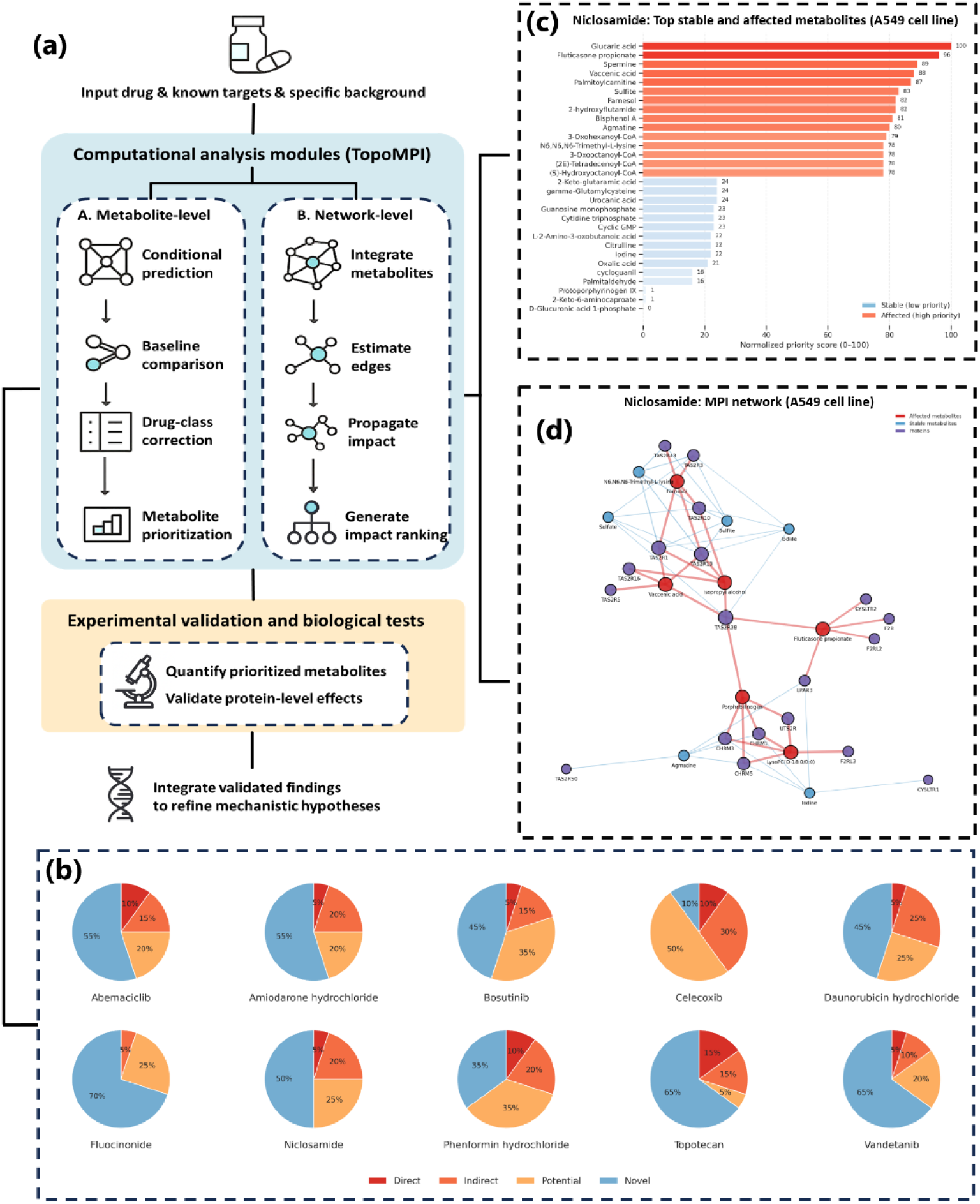
Predicting metabolic impact of drug treatment using TopoMPI. (a) Workflow of the TopoMPI-guided experimental analysis. A schematic overview of the TopoMPI analytical framework integrating computational modeling with experimental design. The pipeline begins with input drug information and known targets, followed by model-based prediction using TopoMPI modules.Two parallel computational paths capture drug-specific effects at the metabolite level (left) and network level (right). **(b) Literature evidence for prioritized metabolite–drug associations.** Literature evidence for prioritized metabolite–drug associations. For each of ten representative drugs, the top 20 ranked metabolite–drug pairs were evaluated through manual literature curation and classified as having direct evidence, indirect mechanistic evidence, contextual co-occurrence, or no prior evidence (“novel”). **(c) Metabolite-level prioritization for Niclosamide.** Normalized priority scores indicate metabolites predicted to be strongly affected (red) or stable (blue) under Niclosamide treatment. High-priority metabolites reflect coherent perturbations across redox, lipid, and energy-related pathways consistent with Niclosamide’s known actions on mTORC1, OXPHOS, AMPK, and Wnt signaling. **(d) Drug-conditioned metabolite–protein interaction network for Niclosamide.** High-priority metabolites (red nodes) propagate their effects to clusters of GPCR and metabolic regulator proteins (violet nodes), forming a coherent perturbed subnetwork. Stable metabolites (blue nodes) display minimal network influence. Edge width reflects metabolite–protein interaction strength, and node size denotes predicted impact. This integrated network highlights biochemical cascades potentially mediating Niclosamide’s metabolic and signaling effects and offers a mechanistic scaffold for targeted experimental validation.

To assess whether the interaction patterns learned by TopoMPI-C reflect meaningful pharmacological similarity, we extracted all protein–metabolite pairs associated with each drug category and computed their distribution across protein functional pathways. These distributions were encoded as interaction profile vectors, capturing each drug category’s preference for targeting specific biological routes. Pairwise cosine similarity between these vectors was used to construct a model-driven similarity network among drug categories.

Comparison of the TopoMPI-C similarity network with four independent drug similarity matrices from Schuhknecht et al. [47] revealed strong concordance, with the model recovering 65.7% (44/67) of known pharmacological similarity edges (**Figure 5c and Supplementary Figure 8**). Beyond recapitulating established relationships, TopoMPI-C uncovered additional similarity links not detected by any reference metric, suggesting previously unrecognized co-regulatory or mechanistic overlap among drug classes. Consistent with this discovery-oriented view, similarity profiles derived from drug–protein–metabolite connectivity (**Supplementary Figure 9**) highlighted coherent clustering of drug categories and biologically related pathway modules, including tight convergence among groups sharing signaling or immune regulatory roles. Together, these findings indicate that TopoMPI-C captures both known and novel dimensions of pharmacological organization, revealing latent functional structure within drug-induced metabolic regulation.

TopoMPI-C recapitulated biologically meaningful drug–protein–metabolite regulatory patterns, with several predictions aligning closely with established pharmacology. The neuroactive ligand–receptor interaction pathway showed the strongest cumulative flow in the model, driven by OPRK1, which linked CNS, infectious disease, and cardiovascular drug classes. This convergence is consistent with extensive evidence demonstrating OPRK1 modulation by exogenous and endogenous opioids in the CNS [79–81], its involvement in viral replication and neuroinflammatory responses [82,83], and its cardioprotective effects through PI3K/AKT and STAT3/OPA1 signaling in cardiac tissue [84–86].

At the metabolite level, TopoMPI-C revealed robust associations between major therapeutic drug classes and amino acid–related metabolites, a pattern strongly supported by prior studies. CNS-active drugs showed well-characterized metabolic signatures, such as gabapentin-induced elevation of GABA [87], levodopa-associated changes in tyrosine, glycine, and homocysteine [88,89], and the broad amino acid reductions induced by tricyclic antidepressants [90–92]. Anti-infective predictions were likewise validated: rifampicin perturbs glycine and arginine metabolism [93], ciprofloxacin disrupts nitrogen-related pathways [94,95], and clarithromycin modulates phenylalanine, tyrosine, and serine metabolism [96,97]. Cardiovascular drug predictions also aligned with known mechanisms, including ACE inhibitor– and ARB-mediated angiotensin metabolism linked to amino acid pathways [98–100], losartan-driven restoration of glutamate derivatives [101], and simvastatin-associated shifts in phenylalanine, histidine, and threonine [102–104]. Together, these observations demonstrate that TopoMPI-C not only captures established pharmacometabolic effects but also highlights underexplored regulatory relationships, revealing latent drug–function connections in metabolic networks.

Histone-modifying enzymes provide a biologically coherent testbed for evaluating how TopoMPI integrates direct metabolic interactions, higher-order associations, and drug impact on metabolism. Histone acetylation, deacetylation, methylation and demethylation act as metabolic– epigenetic interfaces [105], linking cellular metabolic states to transcriptional control and disease processes [106–109]. Across 56 histone-related proteins spanning four enzyme classes, TopoMPI jointly identified 21 proteins connected to 293 metabolites and 46 drug perturbations (**Supplementary Table 5**), offering an integrated view of metabolic influences on chromatin regulation. Across methylases, demethylases, deacetylases, and acetylases (**Figure 5d and Supplementary Figure 10a – c**), TopoMPI consistently identified mechanistically coherent metabolic influences on chromatin enzymes. For methylases, the model linked sterol metabolism with ubiquinone-linked methyltransferase activity through predictions such as COQ5–cholesterol and COQ5–QH₂ [110–112], while also uncovering redox-active polyphenols and oxidative drug conditions modulating TFB2M [113–116]. Demethylase predictions extended these themes, positioning JMJD6 within sterol-regulated pathways [117–119], identifying polyphenol-mediated inhibition of KDM6B [120,121], and highlighting autophagy – lipid regulatory axes via KDM2B – 3-methyladenine –lomitapide [122,123]. Consensus edges emphasized core dioxygenase dependencies on oxygen and mitochondrial redox ratios (KDM4A–oxygen; KDM6B–ubiquinone-2 [124–127]). For acetylases, TopoMPI linked KAT2B to sterols and neurotransmitters [128–130] and revealed drug-responsive redox modulation of PIGL via γ -glutamylcysteine and prifinium bromide [131]. Consensus predictions across all models captured conserved biochemical sensitivities governing deacetylase function, including aldehyde-driven chromatin disruption (HDAC2 – acetaldehyde [132,133]), calcium-channel – linked remodeling (HDAC – nifedipine [134]), and redox modulation by glucosides (HDAC3 – hexyl glucoside [135]). Collectively, these results show that TopoMPI uncovers coherent regulatory logic across histone-modifying enzymes, capturing direct metabolic substrates, redox-driven control, and drug-conditioned signaling that jointly shape chromatin states.

#### 2.4.2 Integrative insights into metabolic effects of drug treatment

By integrating outputs from all three submodels, the framework prioritizes metabolites and proteins most likely to be perturbed under a given drug treatment (**Figure 6a**). Drug-conditioned metabolite predictions from TopoMPI-C are contrasted with baseline estimates from TopoMPI-D and TopoMPI-I to quantify drug-induced deviations and class-specific relevance. These signals are integrated into a composite prioritization score that ranks metabolites according to predicted sensitivity or stability under drug exposure. High-priority metabolites are then mapped onto the baseline metabolite–protein interaction network to infer protein-level propagation patterns, yielding a ranked set of candidate proteins that may mediate downstream biochemical effects.

To evaluate the biological relevance of the prioritization framework, we performed literature-based assessment across 10 representative drugs that are frequently used clinically, examining the top 20 ranked metabolite–drug pairs for each compound. Many high-ranking associations were supported by existing evidence spanning direct metabolite modulation, mechanistic or pathway-level connections, or contextual co-occurrence [136–228], while the remaining predictions represent plausible but previously unreported candidates (**Figure 6b; Supplementary Table 6**). Applying this framework to Niclosamide revealed a metabolic signature that aligns closely with its known actions on mTORC1 inhibition, oxidative phosphorylation (OXPHOS) uncoupling, AMPK activation, and Wnt/β-catenin suppression [137–144]. High-priority metabolites such as Glucaric acid, Vaccenic acid, Palmitoylcarnitine and Spermine (**Figure 6c**) converge on redox regulation, lipid remodeling, fatty-acid oxidation, and polyamine-dependent signaling, pathways previously implicated in Niclosamide’s mitochondrial and energy-stress responses [137,138]. When projected onto the MPI network (**Figure 6d**), these metabolites formed a coherent perturbed subnetwork that propagated strongly to GPCR and metabolic regulator families, including TAS2R, CHRM, and UTS2R, suggesting receptor-linked signaling as a potential route through which Niclosamide transmits metabolic stress signals. In contrast, low-priority metabolites such as Urocanic acid, Cytidine triphosphate, and Oxalic acid remained topologically stable, providing a reference set for normalization and specificity controls. Collectively, these results demonstrate how TopoMPI infers a metabolic landscape upon drug perturbations and uncovers new metabolite–protein interactions that may mediate a drug’s biochemical effects.

## 3. Discussion

In this work, we introduce TopoMPI, a unified heterogeneous graph learning framework that integrates metabolites, proteins, and drugs into a single representational space for systematic inference of metabolite–protein interactions (MPIs). By decomposing the problem into direct interaction prediction (TopoMPI-D), higher-order association modeling (TopoMPI-I), and drug-conditioned regulation (TopoMPI-C), the framework captures complementary layers of metabolic regulation. Together, these modules provide a multi-resolution view of MPI architecture across tissues, perturbational contexts, and molecular modalities.

TopoMPI-D demonstrated strong capacity to recover missing direct MPIs and substantially improved the structural and functional coherence of metabolic networks across 24 tissues. Prediction-augmented networks showed enhanced pathway connectivity and increased centrality of biologically validated nodes, indicating preferential reconstruction of core metabolic regulatory structures. TopoMPI-I expanded this view by identifying topologically indirect but functionally cohesive associations. Its predictions mapped onto enriched protein–protein interaction modules and exhibited stronger diffusion potential toward regulatory hubs, suggesting that indirect learning captures latent biochemical coupling not observable from explicit edges alone. TopoMPI-C further extended the framework by modeling drug-induced rewiring. The resulting tripartite drug–protein–metabolite maps recovered the majority of known pharmacological relationships and revealed new candidate co-regulatory mechanisms at the interface of metabolic and signaling pathways. Collectively, these results show that integrating direct, indirect, and conditional modeling enables TopoMPI to reconstruct biologically meaningful metabolic structure at multiple scales.

Despite its performance, several limitations highlight opportunities for future development. MPI datasets remain sparse and largely tissue-agnostic, constraining the resolution of both supervised and topological signals. The current framework also operates on static graphs and therefore cannot capture the dose-, time-, or state-dependent dynamics that characterize metabolic regulation, particularly under drug perturbation. Moreover, while pretrained embeddings enrich molecular representation, they lack explicit biophysical context such as protein conformations, reaction energetics, or metabolic flux constraints. The heterogeneous edge types in TopoMPI-I and TopoMPI-C are not yet semantically disentangled, and TopoMPI-C does not explicitly encode regulatory directionality or sequential progression of drug–protein–metabolite events. In the future, integrating single-cell multi-omics and time-series drug-response profiles could enable TopoMPI to capture temporal and context-specific metabolic regulation. Structural refinement of heterogeneous message passing, incorporation of causal priors, and explicit modeling of biochemical directionality may further enhance interpretability for pharmacological mechanism discovery. In sum, TopoMPI provides a foundation for reconstructing MPI networks in diverse biological contexts and may facilitate applications in metabolic engineering, drug target discovery, and mechanistic studies of metabolic disease.

## 4. Methods

### 4.1 Data Sources

To construct the heterogeneous interaction graph for the TopoMPI framework, we integrated multiple biological databases and condition-specific datasets, as described below:

(1) Base heterogeneous network construction

The TopoMPI-D model operates on a foundational heterogeneous graph incorporating three types of biological relationships:

Metabolite–metabolite interactions (MMIs) were extracted from co-occurring metabolite pairs within individual reactions in the Recon3D model [44], yielding 16,464 edges.

Protein–protein interactions (PPIs) were obtained from the STRING database [45] and filtered for combined scores ≥ 500, resulting in 51,088 high-confidence edges. The combined score, as defined in the STRING database, is calculated by integrating probabilities from multiple evidence channels and correcting for the chance of random co-occurrence; its values range from 0 to 999, with higher scores indicating stronger confidence in the interaction.

Metabolite–protein interactions (MPIs) were derived from the STITCH database [46] under the same score threshold, retaining 53,392 edges.

The final network comprises 2,375 metabolite nodes and 6,027 protein nodes. MPI edges with scores ≥ 900 were used as positive samples in the training set, and those within the range (500, 900) served as high-confidence test samples.

(2) Context-specific modeling extension

To model condition-specific regulatory patterns, TopoMPI-I and TopoMPI-C incorporate metabolomics data from the A549 cell line (a KRAS-mutant human lung adenocarcinoma model), as reported by Schuhknecht et al. [47]. In that study, the authors developed a high-throughput platform that combines time-lapse microscopy with untargeted metabolomics to profile drug-induced metabolic responses. The dataset includes 1,503 drug perturbations and relative abundance changes in over 1,800 metabolites, enabling system-level analysis of compound-specific regulatory effects.

From this dataset, we extracted a total of 2,082 high-confidence triplets—each representing a significant association between a drug, a target protein (gene), and a downstream metabolite—to serve as positive training instances. In addition, we generated 150,044 negative samples from non-significant associations to create a balanced training set for learning drug-dependent regulatory relationships.:

TopoMPI-I constructs an indirect MPI dataset by extracting correlations between metabolites and proteins across the entire dataset, capturing associations between functionally relevant but topologically unconnected node pairs.

TopoMPI-C further integrates drug perturbation conditions, adding drug–protein interactions (DPIs) and drug–drug structural similarity (DDIs) to form a tri-modal heterogeneous graph, thereby modeling regulatory pathways under pharmacological perturbations.

(3) Tissue-specific network construction

As described in Section 2.2.1, we constructed 24 tissue-specific MPI networks to evaluate the adaptability of TopoMPI-D across diverse tissue backgrounds. The underlying tissue-specific PPIs were curated by Li et al. [50], who processed single-cell omics data from the Tabula Sapiens project [49] to infer protein–protein interaction networks specific to 24 major human tissues.

Building on their results, we constructed tissue-specific heterogeneous networks by filtering our global MPI–PPI–MMI graph according to each tissue’s PPI edge list. For each tissue, only those metabolite–protein interactions and metabolite–metabolite interactions that involved proteins retained in the corresponding tissue-specific PPI network were included. This approach ensures that each tissue network retains consistent graph schema while reflecting biologically plausible protein contexts. We then applied the TopoMPI-D model independently to each of the 24 filtered networks to generate tissue-specific MPI predictions.

(4) Functional validation in blood environment

To validate the biological plausibility of model predictions in physiologically relevant settings (Sections 2.2.2 and 2.3.2), we used a curated set of high-confidence metabolite–protein associations experimentally identified in human plasma by Benson et al. [67]. This dataset served as an independent benchmark for evaluating prediction validity in the blood context.

(5) External PPI networks for indirect interaction validation

In Section 2.3.1, we employed tumor-derived, tissue-specific PPI networks published by Laman Trip et al. [68] as an independent validation source for TopoMPI-I. To ensure contextual alignment with our base networks, we selected four tissues—blood, lung, liver, and kidney—that directly correspond to the 24-tissue network subset, assessing whether TopoMPI-I predictions reflect biologically plausible protein association.

(6) Validation of drug-level regulatory similarity

In Section 2.4.2, we assessed the biological plausibility of drug-level regulatory patterns predicted by TopoMPI-C using four similarity matrices provided by Schuhknecht et al. [47], encompassing iSim[229], chemical structure, mutual information, and Spearman correlation. These external references were used to benchmark the TopoMPI-C–inferred drug similarity network, evaluating its ability to recover known or plausible regulatory commonalities. Notably, the similarity data were sourced from the same dataset used to construct the model input, ensuring consistency across input and evaluation.

### 4.2 TopoMPI Architecture

The TopoMPI framework includes three heterogeneous graph neural network (GNN) models—TopoMPI-D, TopoMPI-I, and TopoMPI-C—each implemented to capture specific types of MPI patterns across biological contexts.

**TopoMPI-D:** This model operates on a static heterogeneous graph comprising MMI, PPI, and MPI edges. Node features are first projected into a common latent space, followed by two GATConv layers [230] that apply relation-aware attention over neighboring nodes. For each metabolite–protein pair, bidirectional embeddings are concatenated and passed through a non-linear MLP classifier to predict interaction likelihood. The model is optimized for binary classification of direct edges and focuses on extracting semantic structure from local neighborhoods.

**TopoMPI-I:** To model topologically unconnected but functionally associated MPI pairs, TopoMPI-I replaces the attention-based GATConv layers with SAGEConv layers [231], which aggregate neighbor information uniformly, expanding the receptive field. It also incorporates a Jumping Knowledge (JK) mechanism [232] to merge representations from different convolution layers, enabling the capture of multi-scale semantic dependencies. Pairwise similarity between metabolite and protein embeddings is used to estimate association strength.

**TopoMPI-C:** Extending TopoMPI-I to a triplet setting, TopoMPI-C introduces drug nodes and two additional edge types: drug–protein interactions (DPIs) and drug–drug similarities (DDIs), forming a tri-modal graph. Separate projection channels are created for drugs, proteins, and metabolites. The model encodes each node type using SAGEConv layers and fuses triplet representations using a shared embedding fusion module, followed by a triplet-level MLP classifier to predict whether a given drug–protein–metabolite unit forms a regulatory interaction under perturbation.

#### 4.2.1 Task Definitions

TopoMPI-D formulates direct MPI prediction as a binary edge classification task on a heterogeneous graph *G* = (*V*, *E*) , where the node set *V* = *V*_*m*_ ∪ *V*_*p*_ includes metabolites and proteins, and the edge set EEE comprises MMI (metabolite–metabolite), PPI (protein–protein), and MPI (metabolite–protein) interactions. The goal is to learn a scoring function:

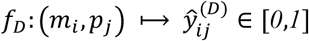

where 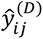 represents the probability that a direct MPI edge exists between (*m* , *p* ). Positive training samples are drawn from high-confidence STITCH interactions, while negatives are randomly sampled unconnected pairs.

TopoMPI-I defines the task as latent association inference between topologically unconnected metabolite–protein pairs. Rather than predicting missing edges, the model estimates a similarity score in the representation space via a function:

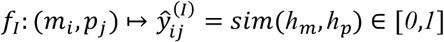

Where *ℎ*_*m*_, *ℎ*_*p*_are node embeddings and 𝑠𝑖*m*(⋅) is a learned nonlinear similarity function. Positive samples are derived from expression-based co-associations, while negatives are weakly or uncorrelated pairs. The predicted scores reflect structural accessibility or potential functional coupling rather than physical interaction.

**TopoMPI-C** further extends the prediction space to conditional triplets (𝑑_𝑘_, *p*_𝑗_, *m*_𝑖_), incorporating drugs dkd_kdk as perturbational context. The objective is to learn:

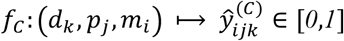

The function indicates the likelihood that *p*_𝑗_mediates a regulatory response on *m*_𝑖_under drug 𝑑_𝑘_. The model encodes DPI (drug–protein), DDI (drug–drug), and MPI relationships to jointly learn regulatory cascades. Training triplets are derived from experimentally observed drug-induced metabolic shifts.

#### 4.2.2 Model Architectures

TopoMPI-D is constructed based on a heterogeneous graph attention network (Hetero-GAT), specifically designed to model semantic dependencies associated with explicit interaction edges. The input graph comprises protein and metabolite nodes, with edge types including MPIs (target edges), PPIs, and MMIs (supporting structure edges). The model architecture consists of the following components:

(1) **Linear Projection Layers**: Metabolite (768-dim) and protein (1280-dim) input features are first mapped into a shared latent space to enable unified convolutional processing:

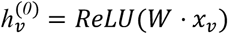

where 𝑥_𝑣_ is the original feature of node 𝑣, and 𝑊 is the learnable projection matrix.

(2) **Two-layer Heterogeneous Graph Convolution**: The model uses two HeteroConv layers, each employing independent GATConv modules for each edge type. Attention-based message passing is applied per relation:

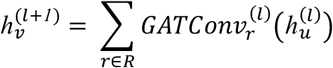

where 𝑅 = {(*p*𝑟𝑜𝑡𝑒𝑖𝑛, *p*𝑟𝑜𝑡𝑒𝑖𝑛), (*m*𝑒𝑡𝑎𝑏𝑜𝑙𝑖𝑡𝑒, *m*𝑒𝑡𝑎𝑏𝑜𝑙𝑖𝑡𝑒), (*m*𝑒𝑡𝑎𝑏𝑜𝑙𝑖𝑡𝑒, *p*𝑟𝑜𝑡𝑒𝑖𝑛)} , 𝑢 is a neighbor of 𝑣, and 𝑙 is the layer index.

(3) **Edge-level Prediction Module:** Final embeddings of metabolite–protein pairs are concatenated and passed through a multi-layer perceptron (MLP) to estimate interaction probability:

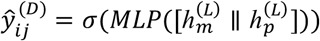

(4) **Loss Function**:To address severe class imbalance, we adopt the Focal Loss:

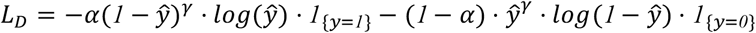

where 𝑦 ∈ {*0*,*1*} is the ground truth label, 𝑦^ is the model output, 𝛼 adjusts class weight, and 𝛾 ≥ *0* controls focus on hard examples.

TopoMPI-I builds on the TopoMPI-D structure with key modifications to support the identification of functionally coupled, but unlinked, metabolite–protein pairs. Its design enhances structural expressiveness and topological abstraction:

(1) **Graph Convolution Replacement**: All GATConv layers are replaced with SAGEConv, better suited for sparse high-order contexts:

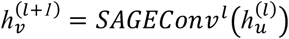

(2) **Jumping Knowledge Aggregation**: To capture multi-scale topological semantics, a JK module aggregates representations from all layers:

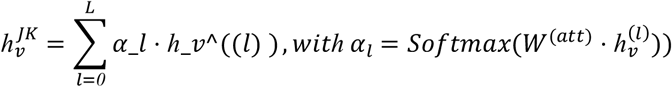

(3) **Pairwise Similarity Inference**: The final metabolite–protein score is obtained by feeding concatenated JK embeddings into an MLP:

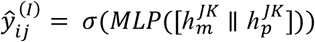

(4) Loss Function: Focal Loss is again used to ensure robustness in sparse positive sample regimes:

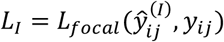

TopoMPI-C extends TopoMPI-I to a tri-modal architecture that incorporates drug nodes to model regulatory mechanisms under drug perturbation. It processes five edge types (MPI, MMI, PPI, DPI, DDI) and performs joint embedding of drug, protein, and metabolite nodes:

(1) **Three-way Feature Projection**: Raw features for drugs (768-dim), metabolites (768-dim), and proteins (1280-dim) are each mapped into a shared hidden space:

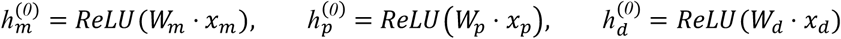

(2) **Multi-relational Heterogeneous Graph Convolution**: Five types of SAGEConv modules are used to update node states per relation, capturing diverse biological dependencies
(3) **Cross-layer Fusion & Triplet Integration**: JK-aggregated embeddings from the three node types are concatenated and fed to an MLP for triplet classification:

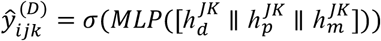

(4) **Loss Function**: Focal Loss is used to address triplet sparsity and enhance training stability:

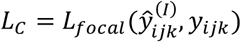

### 4.3 Robustness and Structural Dependency Analysis

**4.3.1 Fast Gradient Sign Method (FGSM) Adversarial Perturbation**

The Fast Gradient Sign Method (FGSM) is a classic one-step adversarial attack technique that generates perturbations in the direction most likely to mislead the model, based on the gradient of the loss function with respect to the input features. For each node *v*, the perturbed feature is computed as:

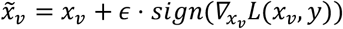

where 𝑥_𝑣_is the original input feature of node 𝑣, 𝐿 is the binary cross-entropy loss used during training, and 𝜖 is a tunable hyperparameter controlling perturbation strength. The perturbation is applied only to the input features, leaving the graph structure and model parameters unchanged.

In our experiments, the perturbations are applied post-training with model weights frozen. We calculate the gradient direction for metabolite and protein nodes in the validation and test sets, and construct corresponding perturbed feature inputs. The experiments are conducted with perturbation strengths 𝜖 ∈ {*0*,*0*.*01*,*0*.*05*,*0*.*1*,*0*.*2*} , covering a spectrum from no perturbation to substantial adversarial noise. For each value of 𝜖, performance metrics (e.g., AUC, accuracy, F1 score) before and after perturbation are recorded to assess model sensitivity to minimal adversarial changes and the stability of its decision boundaries.

#### 4.3.2 Gaussian Noise Injection

Gaussian noise injection simulates random perturbations that arise in real biological data due to measurement noise, incomplete sampling, or nonsystematic variation. Perturbation is defined as:

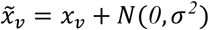

where 𝑁(*0*, 𝜎*^2^*) is a zero-mean Gaussian distribution with variance 𝜎*^2^*, controlling the perturbation magnitude.

Similar to FGSM, noise is applied during testing on fully trained models with fixed parameters and graph structure. For each noise level 𝜎 ∈ {*0*,*0*.*01*,*0*.*05*,*0*.*1*,*0*.*2*}, we independently perturb the features of all metabolite and protein nodes in the test set. Each noise condition is repeated three independent runs, and performance metrics are averaged to mitigate stochastic variation. The perturbed features are used to re-run inference, and changes in performance are compared against the unperturbed baseline to evaluate the model’s robustness to average-case input fluctuations.

#### 4.3.3 Ablation of Graph Structural Components

To assess the contribution of auxiliary biological networks to MPI prediction, we performed ablation experiments in which specific edge types were selectively removed from the heterogeneous graph prior to model training. Three configurations were evaluated: (1) Removal of PPI edges only; (2) Removal of MMI edges only; (3) Removal of both PPI and MMI edges simultaneously.

For each configuration, all three TopoMPI submodels were retrained using identical hyperparameters, training–validation splits, and node feature embeddings. Performance differences relative to the full graph quantified the extent to which protein–protein connectivity and metabolite–metabolite similarity support the structural reasoning capabilities of each model.

### 4.4 Topological Validation of TopoMPI-D

To systematically assess the structural enhancement capabilities of TopoMPI-D in tissue-specific regulatory networks, we designed a network topology validation framework based on comparisons between an original network and its corresponding extended versions. All networks were constructed using the blood tissue context. The original network, as defined in Section 2.2.1, integrates high-confidence protein–protein interactions (PPIs), metabolite–metabolite interactions (MMIs), and metabolite–protein interactions (MPIs) to represent a baseline biological regulatory map within the blood environment.

To evaluate whether the predicted edges from TopoMPI-D lead to topological improvement, we generated three extended networks by adding model-predicted metabolite–protein edges to the original network. These predictions were not added indiscriminately; instead, edges were selected through a combination of statistical significance testing and score-based ranking. Specifically, all candidate metabolite–protein edges predicted by the model were first collected and subjected to two statistical significance tests: empirical p-values and permutation-based p-values. The empirical p-value was calculated as the proportion of candidate edges with prediction scores greater than or equal to a given edge, while the permutation p-value was derived via Monte Carlo sampling (n = 10,000) from the candidate score distribution to construct a non-parametric null model.

Edges achieving significance (p < 0.01) under either method were retained. These edges were then ranked by their prediction scores, and three progressively augmented networks were built by sequentially adding the top 5,000, 10,000, and 15,000 edges, respectively.

#### 4.4.1 Node Centrality Analysis

To evaluate how the predicted edges affect the structural importance of key regulatory nodes, we conducted a node-level validation based on standard centrality measures. The analysis focused on metabolite and protein nodes included in the validation set, and examined changes in their topological prominence after edge augmentation.

Centrality Measures

For each version of the network (original and the three augmented variants), we calculated three centrality metrics for all nodes: Degree Centrality, Closeness Centrality, and PageRank. Degree Centrality quantifies the number of direct connections, Closeness Centrality measures the average shortest-path distance to all other nodes, and PageRank estimates the global influence of a node via iterative propagation. All metrics were computed on undirected graphs to maintain consistency in structural comparison.

Validation Set and Pairing Strategy

To ensure biological interpretability, we selected metabolite and protein nodes with literature-backed interactions as the validation set. We extracted their centrality scores across all four networks and matched them by node ID, enabling a direct node-to-node comparison across network versions. This pairing approach ensures that the analysis isolates the change in importance for the same node, independent of global network shifts.

Paired Statistical Testing

A one-sided Wilcoxon signed-rank test was used to evaluate whether the centrality scores of validation nodes increased significantly in the augmented networks compared to the original. The null hypothesis was that the added edges do not increase node centrality. For each centrality metric and augmentation level (Top-5000, Top-10000, Top-15000), we conducted paired comparisons. Median differences and p-values were reported for each combination of node type (protein or metabolite) and centrality metric. Statistical significance thresholds were set at 0.05, 0.01, and 0.001 and denoted using standard notation. This approach provided a quantitative assessment of whether TopoMPI-D successfully promotes the structural importance of biologically validated nodes.

#### 4.4.2 Edge Importance Change

To further evaluate the structural relevance of TopoMPI-D predicted edges, we designed a ranking-based validation framework from the perspective of edge-level importance. This analysis assessed whether the model preferentially introduced structurally critical edges—those that enhance network connectivity, bridge modular boundaries, or integrate functional hubs. Specifically, we focused on two categories of structurally important edges: hub–hub edges and bridge edges, and compared their ranks in the original and augmented networks.

Definitions of Edge Categories

We employed two complementary metrics to define potentially critical edges in the network:

Hub–Hub Edges: Edges that connect two nodes with high centrality. First, Degree, Closeness, and PageRank centralities were computed for all nodes. These values were normalized and averaged to obtain a composite centrality score. For each edge, we calculated the average composite score of its two endpoints to derive the hub–hub score and rank the edges accordingly.

Bridge Edges: Edges with high edge betweenness centrality, indicating their involvement in shortest paths and their role in connecting network modules or bottlenecks.

For each version of the network (original and augmented), we computed both scores and generated edge importance rankings based on these metrics.

Rank Difference Computation and Missing Value Strategy

To quantify changes in structural relevance, we calculated the rank differences for each validation edge between the original network and the augmented network (Top-15000). Since many predicted edges were not present in the original network, they lacked baseline rankings. To handle such missing values, we assigned them a conservative default rank equal to the midpoint of the total rank range in the original network.

This imputation strategy assigns a neutral reference point to missing edges—only edges ranked in the top half of the augmented network are considered as having increased importance. This conservative approach avoids overestimating structural improvement and ensures robustness in comparative analysis.

All ranks were normalized to the [0, 1] range. The ΔNormalized Rank (difference in normalized rank) was computed for each edge, serving as the main indicator of edge-level importance change. Validation Set and Statistical Testing

The analysis focused on metabolite–protein edges supported by external databases or literature. These validated edges were treated as positive instances to assess whether TopoMPI-D successfully identifies biologically meaningful regulatory connections. For each edge type (hub–hub and bridge), paired one-sided Wilcoxon signed-rank tests were performed to determine whether the ΔNormalized Rank in the augmented network was significantly higher than in the original network.

Additionally, we stratified the validation edges by metabolite functional category to evaluate whether the model exhibited consistent structural enhancement across different biochemical domains.

#### 4.4.3 Pathway Connectivity Change

To evaluate whether TopoMPI-D enhances pathway-level connectivity at the system level, and whether such topological changes are biologically meaningful, we developed a functional validation framework based on changes in pathway structural integrity. The core idea was to frame the analysis as a binary classification problem: can changes in topological connectivity distinguish between pathways that include validated MPIs and those that do not?

Connectivity Metrics per Pathway

For each network version (original and Top-15000), we extracted pathway subgraphs using KEGG annotations [233]. For each pathway, we computed the following topological connectivity metrics: Reachability: Proportion of node pairs that are connected via at least one path; Largest Connected Component (LCC) Size: Number of nodes in the largest connected component within the pathway subgraph; Global Efficiency: Average inverse shortest-path length among all node pairs, indicating how efficiently information propagates.

For each metric, we calculated the difference between the augmented and original networks (ΔMetric). Only pathways meeting the following criteria were retained for evaluation: (i) at least three nodes across all networks; (ii) at least one edge in the augmented network.

Biological Labeling and Classification Task

To label pathways as biologically functional or not, we mapped literature-validated MPIs to KEGG annotations. Pathways containing at least one validated MPI were labeled as positive; others were negative. Thus, each pathway became a sample with ΔMetric features and a binary label.

AUC-Based Metric Evaluation

For each ΔMetric, we computed the ROC curve and corresponding AUC to evaluate its ability to discriminate between pathways that contain at least one experimentally validated metabolite–protein interaction and those that do not. We further applied bootstrap resampling (1,000 iterations) to estimate the 95% confidence interval (CI) for each AUC. Statistical significance was assessed by checking whether the lower bound of the AUC CI exceeded 0.5, and significance levels were annotated using standard star notation (*, **, ***).

To explore potential category-specific effects, we also conducted stratified ROC analyses across metabolite categories (e.g., carbohydrates, lipids, organic acids), retaining only categories with at least five validated pathways.

Enrichment Testing

To complement the ROC analysis, we conducted enrichment tests using Fisher’s exact test. For each ΔMetric, we defined the top 25% of pathways (highest ΔMetric) as the “enhanced connectivity” group and tested whether these were enriched for validated pathways. Odds ratios and p-values were reported for both global and category-specific analyses.

### 4.5 Network-Based Validation of TopoMPI-I Predictions

To systematically evaluate the structural and biological plausibility of the indirect metabolite–protein associations predicted by TopoMPI-I, we designed a three-part validation framework based on protein–protein interaction (PPI) network analysis. All analyses leveraged the predicted outputs of TopoMPI-D and TopoMPI-I, as well as a reference PPI network derived from curated validation datasets.

Based on this setup, we constructed four distinct sets of protein–protein pairs for comparison: Only-Direct Group: protein pairs that are indirectly linked through shared metabolites solely in TopoMPI-D predictions; Only-Indirect Group: protein pairs indirectly inferred via high-order topological paths unique to TopoMPI-I predictions; Overlap Group: protein pairs co-predicted by both TopoMPI-D and TopoMPI-I; Random Group: background control group with protein pairs sampled to match the degree distribution of the base network, thereby controlling for node degree bias.

#### 4.5.1 Protein–Protein Interaction Confidence Enrichment

This analysis aimed to assess whether the protein pairs inferred from TopoMPI-I predictions represent plausible biological interactions. For this purpose, we referred to an external validation dataset containing pairwise PPI confidence scores derived from integrated experimental and literature evidence, which served as a proxy for interaction potential.

We quantified and compared the PPI confidence scores across the four groups (Only-Indirect, Only-Direct, Overlap, and Random) to determine whether TopoMPI-I preferentially identifies functionally enriched protein relationships.

For each group, we extracted protein pairs with defined interaction scores in the validation dataset and retained them for statistical evaluation. We then conducted two complementary types of analysis:

(1) Statistical Distribution Comparison: Using three classic tests—Student’s t-test, Mann–Whitney U test, and Kolmogorov–Smirnov test—we evaluated whether the interaction score distributions for the Only-Indirect, Only-Direct, and Overlap groups differed significantly from the Random group. This assessed whether model-derived pairs were systematically enriched for higher-confidence biological interactions.
(2) Density Difference Analysis: We constructed kernel density estimation (KDE) curves for each group and computed the pointwise difference between the Random group and the remaining three groups. This analysis highlighted shifts in score density within specific confidence intervals, providing a more detailed picture of enrichment trends at high-confidence ranges.

#### 4.5.2 Local Topological Proximity

To determine whether TopoMPI-I–predicted protein pairs tend to be embedded within structurally cohesive neighborhoods in the external PPI network, we evaluated the local topological characteristics of subnetworks induced by each group.

For each group of protein pairs, we identified all unique protein nodes involved and extracted the induced subgraph from the external PPI network. On each subgraph, we computed six representative network topology metrics:Average Shortest Path Length: the mean path length between all node pairs in the subgraph (defined only for connected graphs);Density: ratio of observed to possible edges, reflecting overall connectivity;Average Clustering Coefficient: average likelihood of triadic closure among each node’s neighbors; Modularity: strength of community structure using a greedy modularity maximization algorithm;Transitivity: global proportion of closed triplets, indicating the extent of hierarchical clustering;Assortativity: the tendency for nodes of similar degree to be connected.

These metrics jointly characterized each group’s local network cohesion, allowing us to assess whether TopoMPI-I tends to identify functionally integrated protein clusters that are structurally localized within the real-world PPI network.

#### 4.5.3 Network Propagation Capability

We implemented a network diffusion–based analysis to quantify the extent to which predicted proteins from each group propagate toward central nodes in tissue-specific PPI networks.

For each tissue, we constructed an undirected PPI network using interactions with a combined confidence score ≥ 0.8. Each protein group—Only-Direct, Only-Indirect, Overlap, and Random—was used as a seed set in a Random Walk with Restart (RWR) simulation. RWR iteratively spreads influence from seed nodes while intermittently restarting, yielding a stable diffusion score distribution over all nodes.

We defined the “functional core” of the network as the top 500 nodes ranked by degree. For each group, we calculated Precision (K = 100, 200, 500), representing the proportion of top-K diffusion-ranked nodes overlapping with the functional core. To control for group size disparities, we normalized input by subsampling to the smallest group size and repeated each simulation five times, reporting the average and standard deviation of Precision across replicates.

### 4.6 Validation of TopoMPI-C Based on Drug Similarity

To evaluate whether TopoMPI-C is capable of capturing biologically meaningful functional similarities between drugs, we constructed a drug–drug similarity network derived from the model’s predictions and quantitatively compared it against a set of multi-perspective drug similarity reference datasets from the same source publication. This comparison aimed to assess the consistency between TopoMPI-C–inferred relationships and known therapeutic class associations.

Construction of the Drug–Drug Similarity Network via Prediction-Based Jaccard Similarity

We first applied statistical filtering to the full set of TopoMPI-C predictions, retaining only metabolite–protein interaction (MPI) edges with empirical or permutation p-values less than 0.01, following the significance filtering procedure described in Section 4.4. For each drug, the predicted MPI edges were ranked by interaction score, and the top 10% were extracted as representative functional associations. These high-confidence edges were used to build a binary matrix capturing drug–metabolite–protein pairings, which effectively encode each drug’s downstream regulatory profile. Pairwise drug–drug similarities were then computed using the Jaccard index, with higher values indicating greater overlap in predicted metabolite–protein targets and, by extension, potential functional resemblance.

Extraction of Multi-Perspective Drug Similarity Reference Networks

We utilized four drug–drug similarity matrices provided in the same reference (see Section 4.1(6)), each derived from a distinct methodological perspective: iSIM: inferred pharmacological similarity via knowledge graph reasoning;Chemical Similarity: based on molecular structure fingerprints;Mutual Information: derived from mutual information scores across large-scale drug response profiles;Spearman Correlation: capturing monotonic similarity in drug response signatures.

From each matrix, we extracted drug pairs identified as significantly similar and analyzed the distribution of these pairs across known therapeutic classes. Based on these, we constructed data-driven therapeutic class similarity networks. The statistical significance of enriched relationships was replicated following the procedures described in the original publication, using hypergeometric testing and Bonferroni correction for multiple hypothesis adjustment.

Comparative Consistency Between Model-Inferred and Reference Similarity Networks

To systematically assess whether TopoMPI-C successfully reconstructs functional similarities between drug classes, we compared the therapeutic class–level similarity matrices derived from TopoMPI-C predictions against each of the four external reference networks. For each reference network, we calculated the proportion of significant therapeutic class relationships that were correctly recovered in the TopoMPI-C–based similarity network. This served as a direct metric of TopoMPI-C’s consistency in modeling drug-related functional associations from a systems-level perspective.

### 4.7 Experimental guidance analysis: drug-conditioned cross-validation and metabolite/network-level impact scoring

To simulate real experimental conditions and identify metabolites and protein interactions most responsive to individual drug perturbations, a leave-one-drug-out (LODO) cross-validation framework was designed as the first stage of the TopoMPI-based experimental guidance analysis.

Rather than serving as a conventional performance benchmark, this LODO procedure was implemented to emulate in silico “drug omission experiments,” enabling the exploration of metabolic response patterns triggered by unseen drug conditions.

By withholding all information related to a specific drug during model training and then reintroducing it as a test case, TopoMPI generates unbiased predictions of how that drug may perturb metabolite–protein relationships within the global metabolic interaction (MPI) network. Subsequent metabolite- and network-level scoring analyses build upon these predictions, quantifying and ranking the degree of metabolic and proteomic perturbation induced by each compound.

#### 4.7.1 Leave-one-drug-out simulation of drug-induced metabolic perturbation

In the LODO design, each drug 𝑑_𝑖_was sequentially excluded from the training data, and all associated triplets (*m*, *p*, 𝑑_𝑖_) —comprising metabolite, protein, and drug interactions—were withheld. The remaining drugs (𝐷 − 1) formed the training set used to retrain the TopoMPI models from scratch.

After model convergence, the excluded drug 𝑑_𝑖_ was reintroduced as a query compound to generate its conditional metabolite–protein interaction scores 𝑆_𝐶_(𝑑_𝑖_, *m*, *p*). This process was repeated for every drug, producing per-drug prediction tables that mimic how the model would behave for an untested compound. Formally:

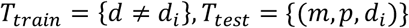

For each LODO iteration, both TopoMPI-D and TopoMPI-I were used to establish baseline, drug-independent M–P association scores 𝑆_𝑏𝑎𝑠𝑒_ (*m*, *p*), while TopoMPI-C produced conditional, drug-contextual predictions 𝑆_𝐶_(𝑑_𝑖_, *m*, *p*).

#### 4.7.2 Metabolite-level impact quantification

At the metabolite layer, TopoMPI-C predictions were combined with baseline outputs from TopoMPI-D/I and drug–drug similarity (DDI) information to quantify the extent to which each metabolite responded to the introduction of a specific drug. Each metabolite *m* under drug 𝑑 was characterized by five complementary quantitative metrics designed to capture different aspects of drug-induced metabolic perturbation.

(1) **conditional confidence score:** 𝑆_𝐶_(𝑑, *m*) represents the direct prediction from TopoMPI-C, quantifying the likelihood that metabolite mmm is affected under the presence of drug *d*. This score reflects the model’s intrinsic confidence in drug-conditioned metabolic regulation.
(2) **empirical significance score:** 𝑞_𝑟𝑎𝑛𝑘_was computed from a rank-based transformation of the conditional scores. Specifically, for each drug 𝑑:

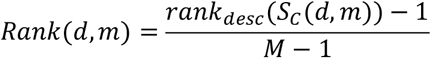

where *M* is the number of metabolites evaluated for *d* and 𝑟𝑎𝑛𝑘_𝑑𝑒𝑠𝑐_ denotes the descending average rank of 𝑆_𝐶_ (ties resolved by average rank). 𝑅𝑎𝑛𝑘(𝑑, *m*) ∈ [0,1] indicates the relative standing of *m* within the drug-specific score distribution (higher values = more extreme/high-confidence effects). We then obtain an empirical p-value as *p*_𝑒*mp*_(𝑑, *m*) = 1 − 𝑅𝑎𝑛𝑘(𝑑, *m*) and apply Benjamini–Hochberg false-discovery rate correction across metabolites to yield 𝑞_𝑟𝑎𝑛𝑘_. This procedure quantifies the statistical significance of each metabolite’s predicted perturbation relative to all other metabolites for the same drug.

(3) **baseline deviation score:** 𝛥(𝑑, *m*) was defined as the difference between the drug-conditioned prediction and the baseline (drug-independent) prediction from TopoMPI-D/I:

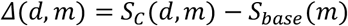

This metric highlights metabolites whose model-predicted interactions deviate strongly from the steady-state network configuration once a drug is introduced, serving as an estimate of metabolic sensitivity to perturbation.

(4) **class-specific specificity score:** 𝑆_*p*𝑒𝑐_ (𝑑, *m*) was computed by comparing each metabolite’s conditional score to the mean prediction across structurally or pharmacologically similar drugs identified in the DDI network:

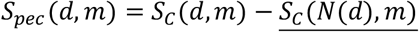

Where 𝑁(𝑑) is the set of DDI-linked neighbor drugs with similarity ≥ 0.8. This measure identifies drug-unique metabolic effects, distinguishing general class-related responses from compound-specific signatures.

(4) **composite priority score**: these four components were integrated into a composite priority score that balances confidence, significance, deviation, and specificity:

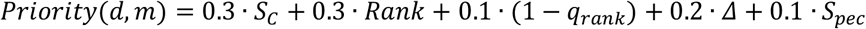

#### 4.7.3 Network-level impact scoring

To translate metabolite-level perturbations into a system-level interpretation, prioritized metabolites were mapped onto the global MPI network to estimate how drug-induced changes may propagate to associated proteins.

(1) **Edge scoring**: Each metabolite–protein edge was assigned a condition-adjusted score:

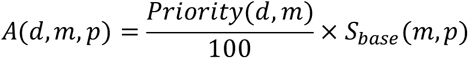

representing the relative impact strength of metabolite *m* on protein *p* in the presence of drug *d*.

(2) **Protein impact scoring**: The cumulative effect of the drug on each protein was then quantified as:

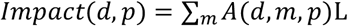

A normalized 𝑖*mp*𝑎𝑐𝑡_𝑛𝑜𝑟*m*_ score (0–100) was used to rank potentially affected proteins.

(3) **Network pruning and visualization**: To maintain interpretability, only the top 5 edges per metabolite were retained, weak baseline edges (𝑆_𝑏𝑎𝑠𝑒_ < 0.3) were excluded, and all known drug targets were preserved.

## Code Availability

The TopoMPI framework was implemented in Python using PyTorch and PyTorch Geometric. All code is available at https://github.com/laxlyt/TopoMPI. The repository includes scripts for model implementation of TopoMPI-D, TopoMPI-I, and TopoMPI-C, as well as example workflows reproducing the main analyses presented in this study. Detailed instructions and configuration files are provided to facilitate reproducibility and reuse.

## Acknowledgments

We gratefully acknowledge support from the National Institutes of Health grant R35 GM13779501 and the Camille and Henry Dreyfus Foundation (to S.C.).

## Author contributions

Conceptualization, S.C, Y.L.; investigation, Y.L.; funding acquisition, S.C.; project administration, S.C.; supervision, S.C.; writing – S.C. and Y.L.

