## Supplementary Figures for "A Heterogeneous Graph Framework for Inference of Metabolite–Protein-Drug Interaction Networks"

**Supplementary Figure 1**
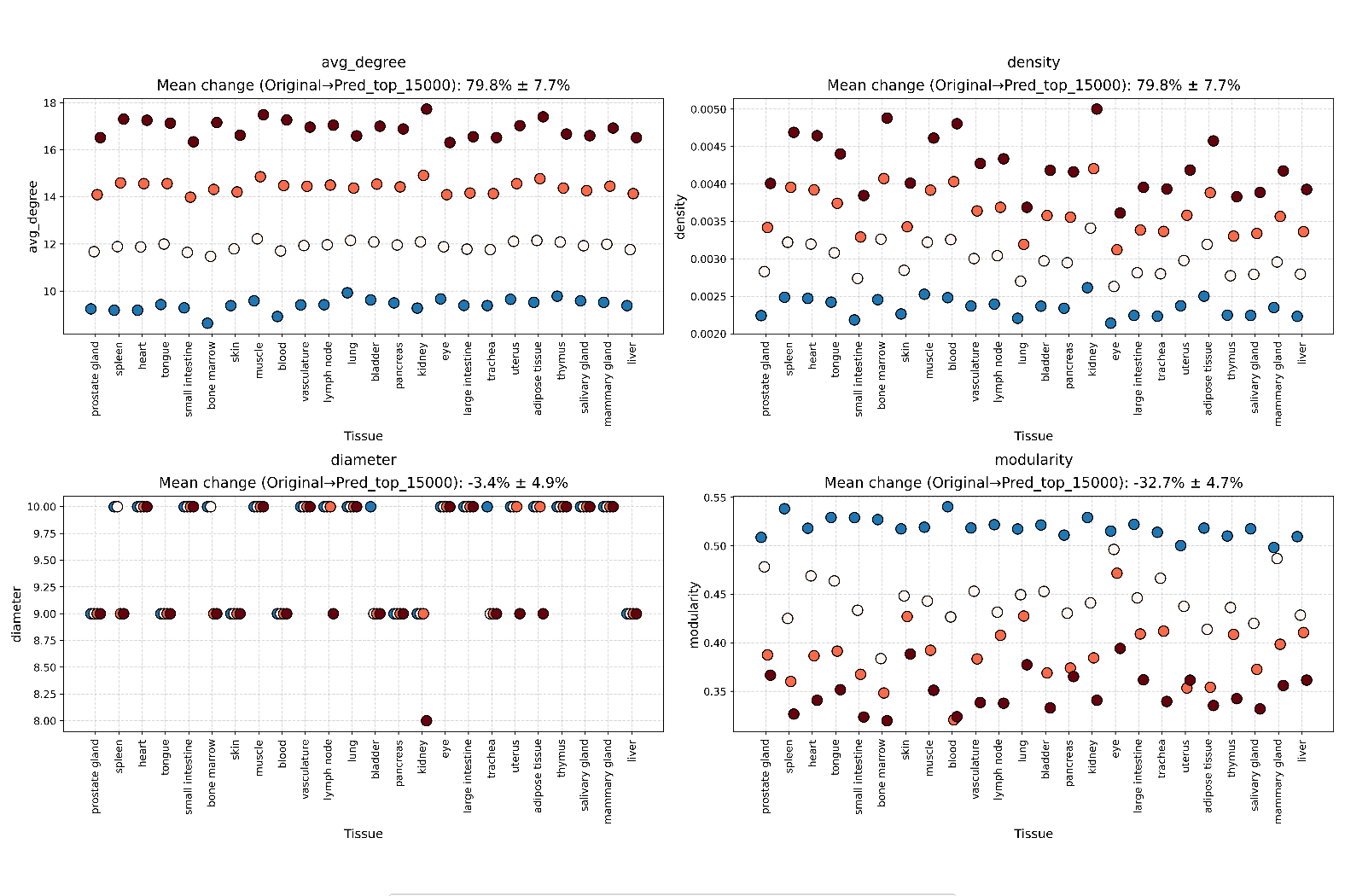

**Comparison of four topological measures across 24 tissue-specific networks before and after edge extension.** From top left to bottom right: average node degree, edge density, network diameter, and modularity. All extended networks exhibited increased connectivity and altered structural properties compared to their original counterparts.

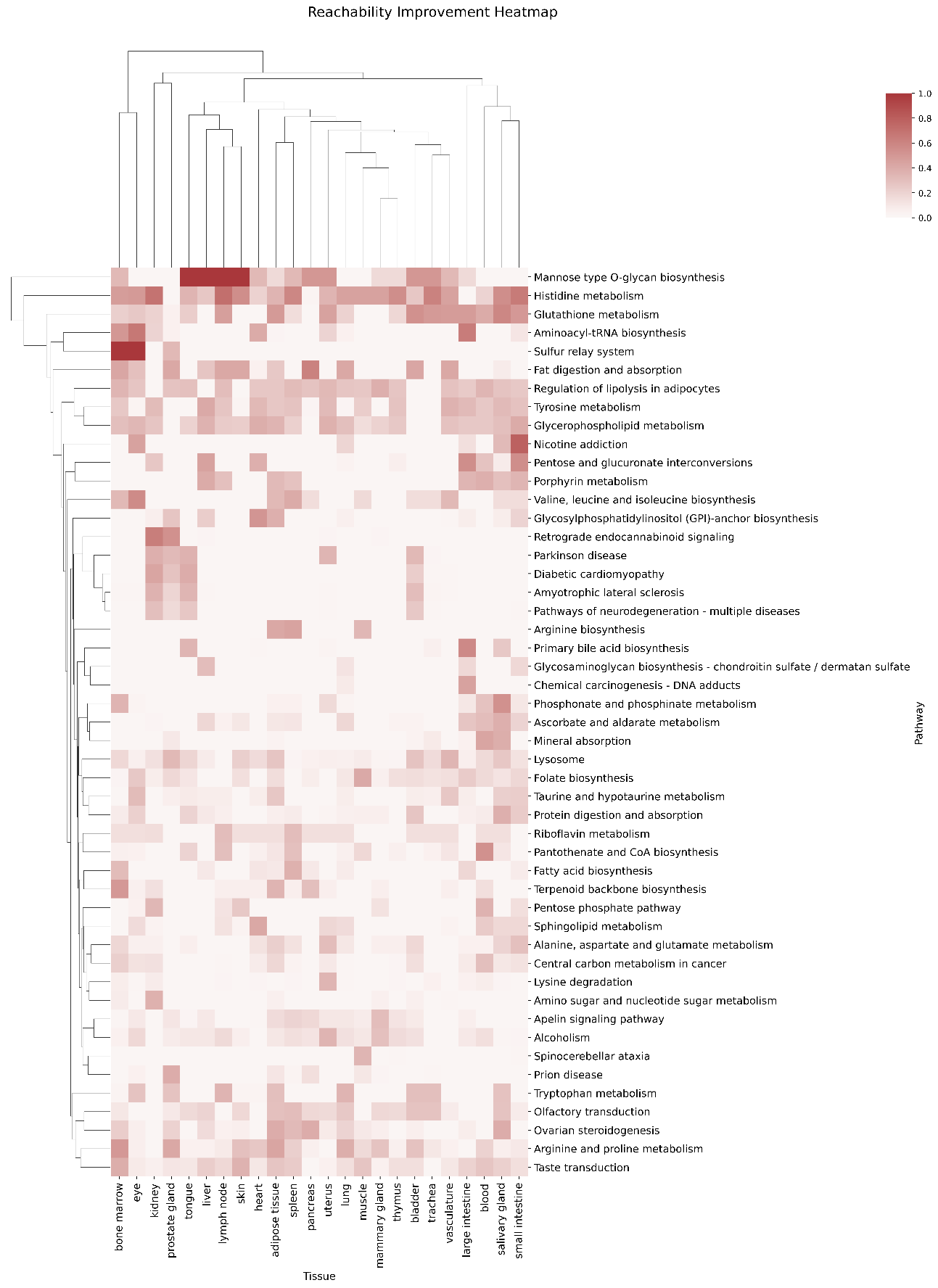
**Supplementary Figure 2**

**Clustering heatmap of changes in pathway connectivity across 24 tissues (Extended vs. Original networks).** For each tissue, we quantified shifts in metabolic pathway connectivity to evaluate whether the predicted MPI edges enhanced structural integration within known pathways. Reachability is the proportion of node pairs that are connected via at least one path. The heatmap reveals heterogeneous gains across tissues, reflecting context-specific improvements in pathway coherence.

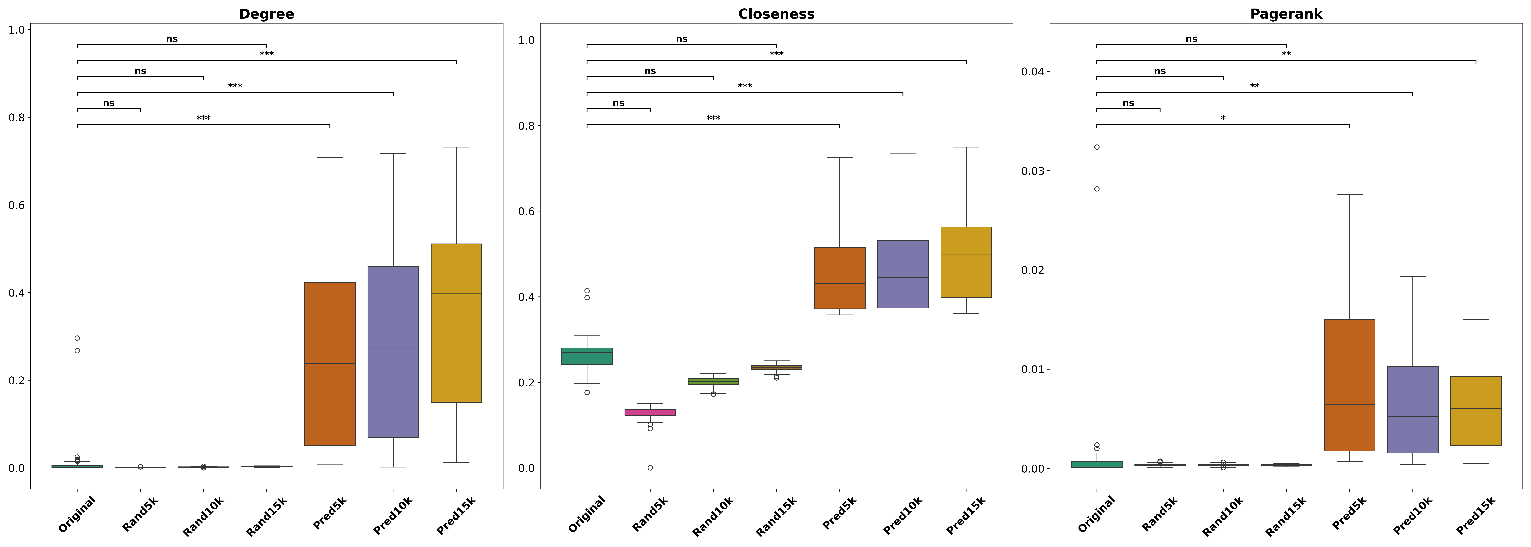
**Supplementary Figure 3**

**Changes in centrality of validation-set metabolite nodes.** This panel presents three centrality metrics—degree, closeness, and PageRank—for experimentally validated protein nodes across four network states (original and three expanded networks with increasing numbers of predicted edges).

**Supplementary Figure 4**
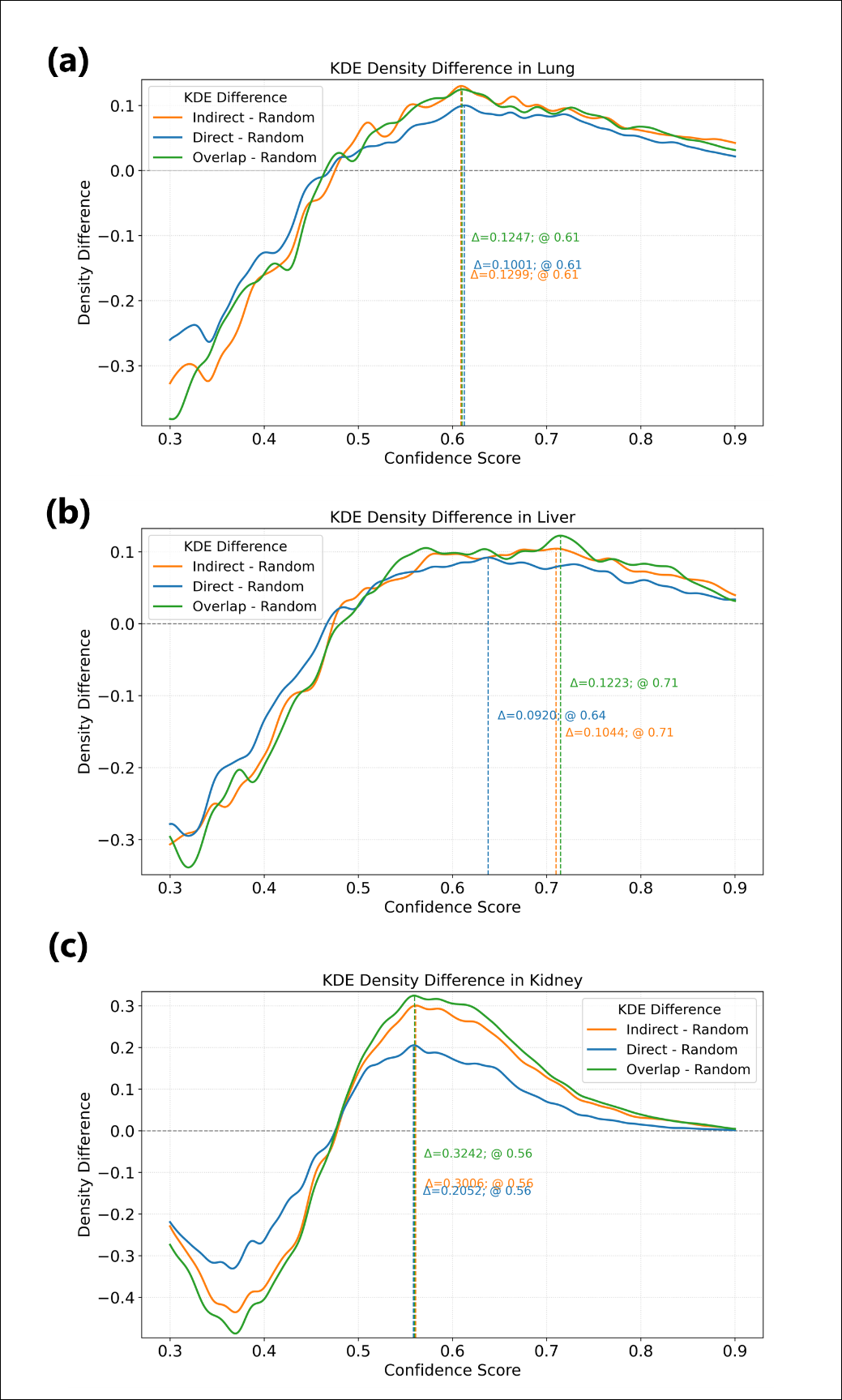

**PPI confidence enrichment in lung, liver and kidney.** Kernel density estimates (KDEs) show the distribution of external confidence scores for protein pairs across four groups. Only-Indirect and Overlap groups exhibit right-shifted density peaks relative to Only-Direct and Random, indicating that TopoMPI-I preferentially recovers high-confidence interactions overlooked by direct predictions.

**Supplementary Figure 5**
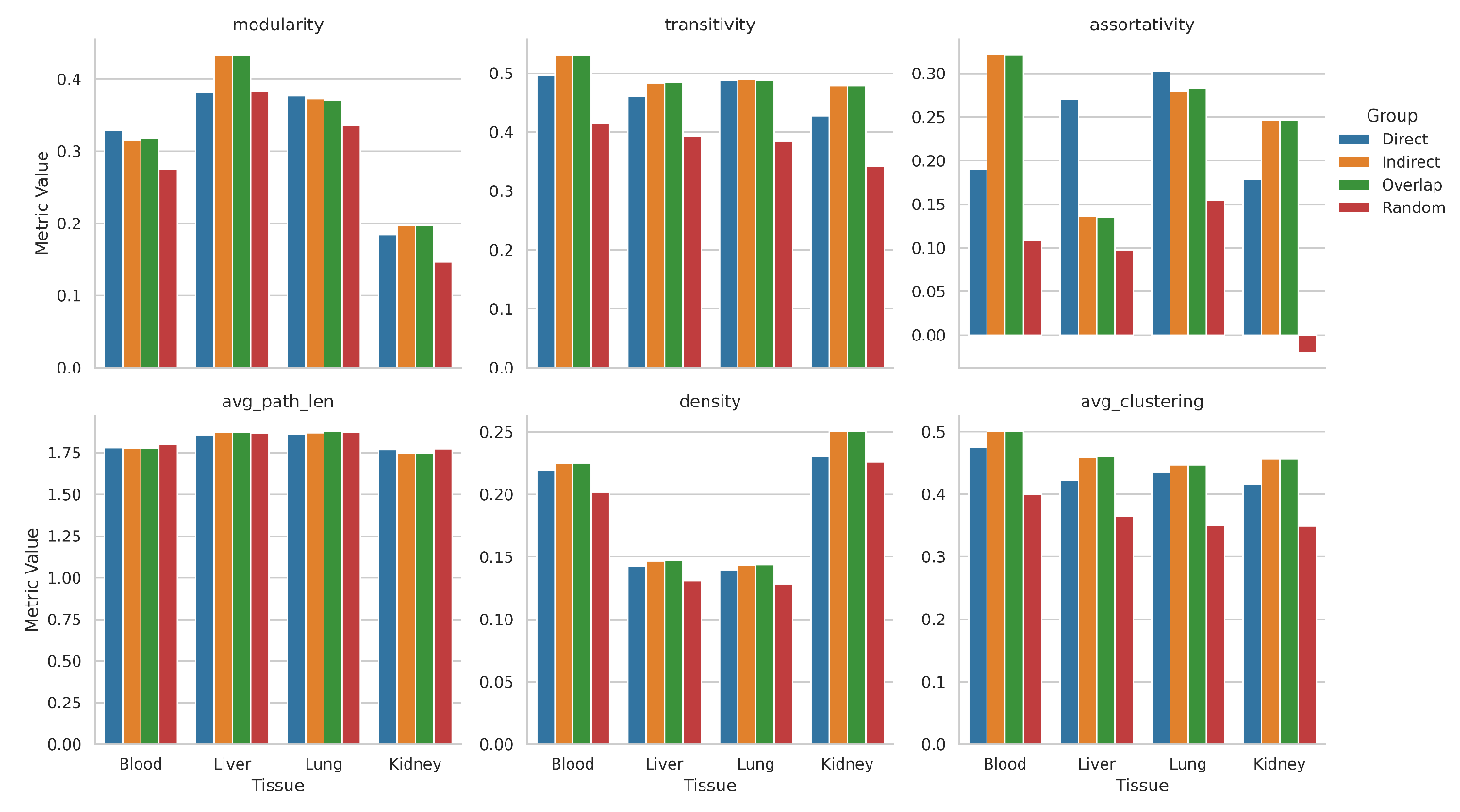

**Topological feature comparison.** For each group, subgraphs were extracted and evaluated using six network metrics: modularity, transitivity, assortativity, average path length, density, and average clustering coefficient. Predicted groups outperform the random group on clustering and modularity metrics, indicating that MPINet-I predictions are concentrated in structurally cohesive and functionally enriched regions.

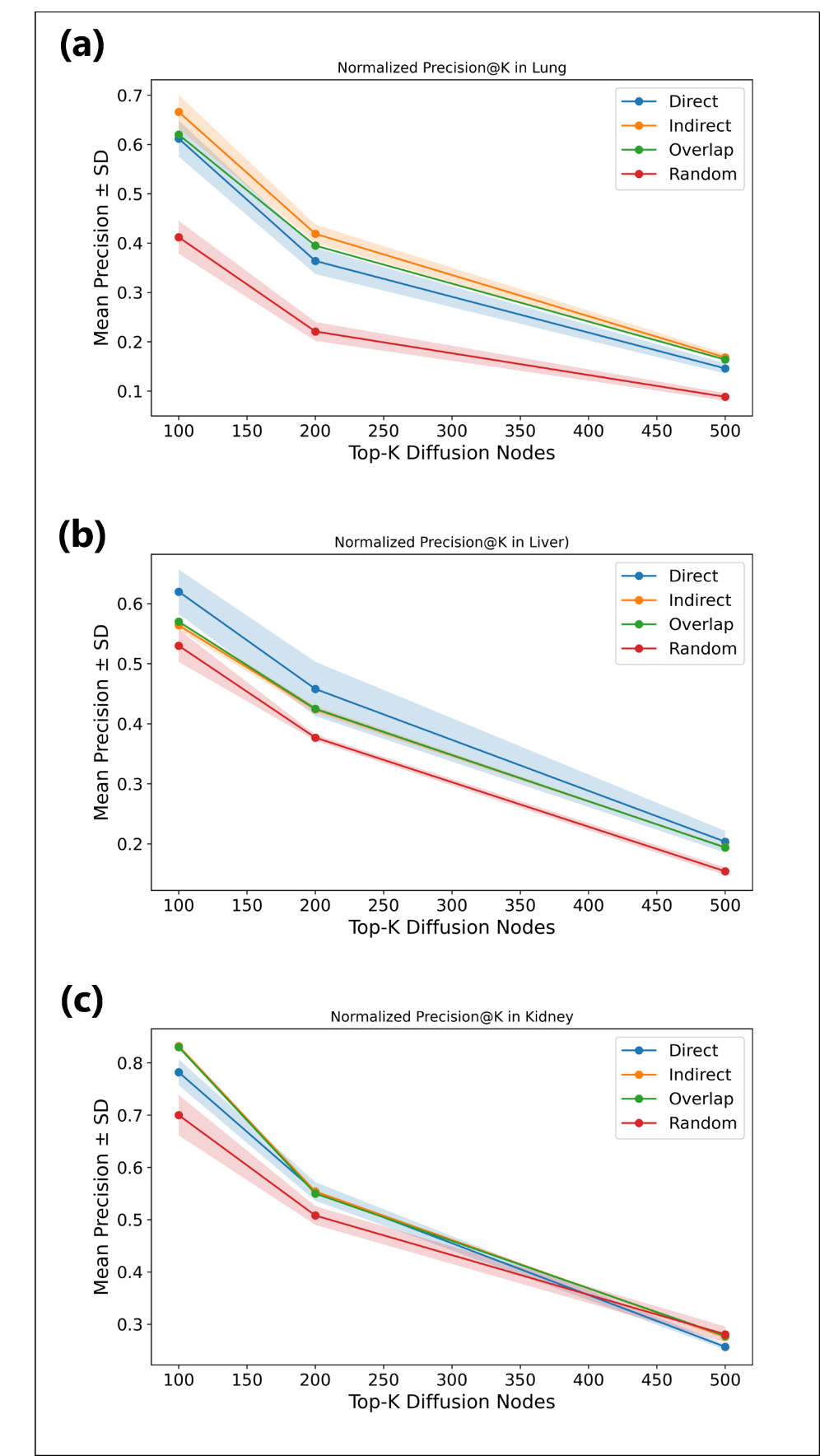
**Supplementary Figure 6**

**Network propagation analysis in lung, liver and kidney.** Using Random Walk with Restart (RWR) on a high-confidence PPI network, Precision (K = 100, 200, 500) is computed based on overlap with top PageRank nodes. Indirect predictions consistently achieve higher Precision（K = 100 and 200) than Direct-only, indicating stronger embedding in core regulatory regions.

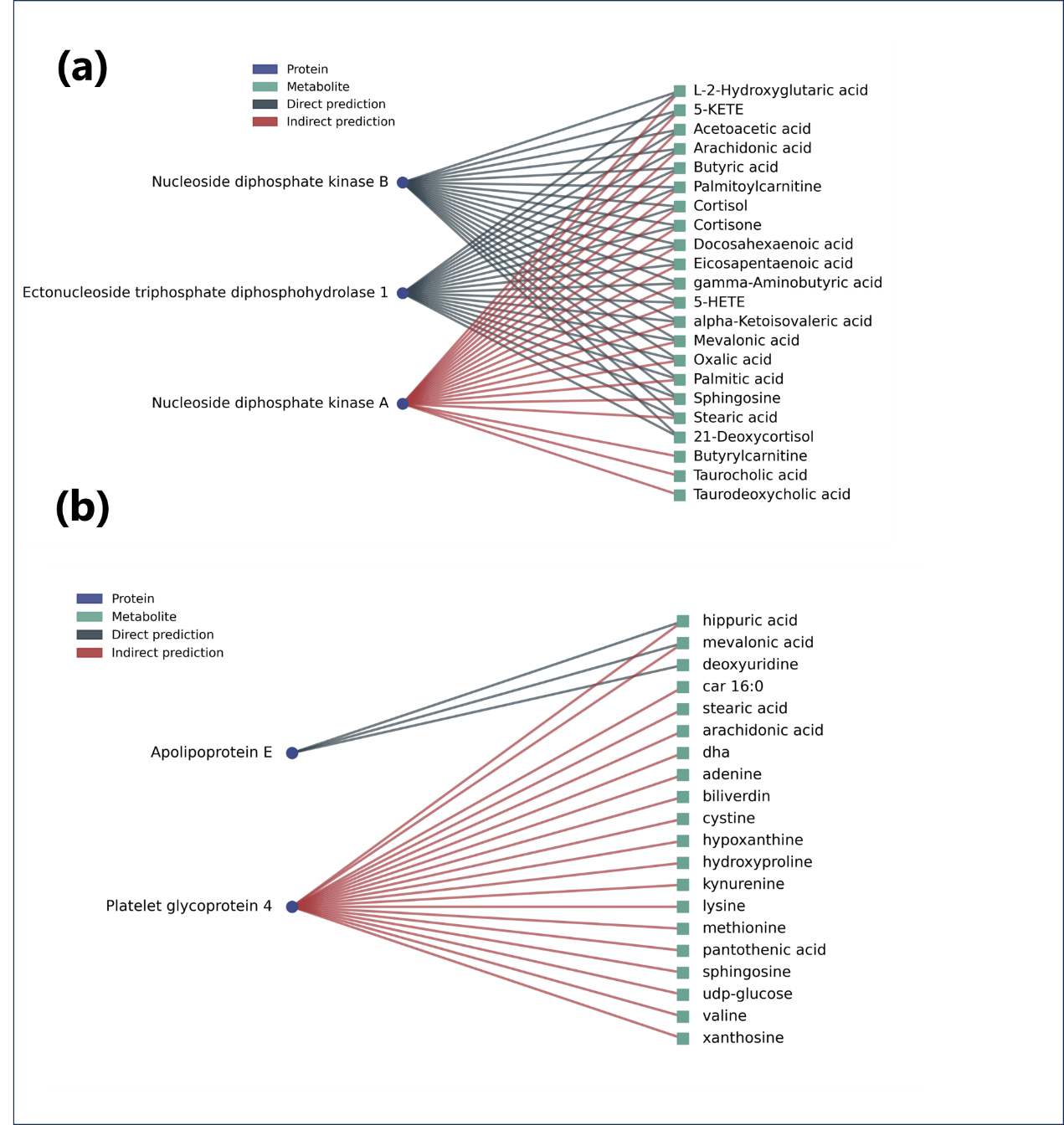
**Supplementary Figure 7**

**Case studies in the blood tissue network.** Representative Only-Indirect predictions are shown in three functional contexts: (a) nucleotide metabolism; (b) a regulatory module centered on APOE and CD36. These interactions are absent from MPINet-D predictions but are supported by prior literature, demonstrating that MPINet-I complements direct predictions by uncovering biologically validated but structurally non-obvious interactions.

**Supplementary Figure 8**

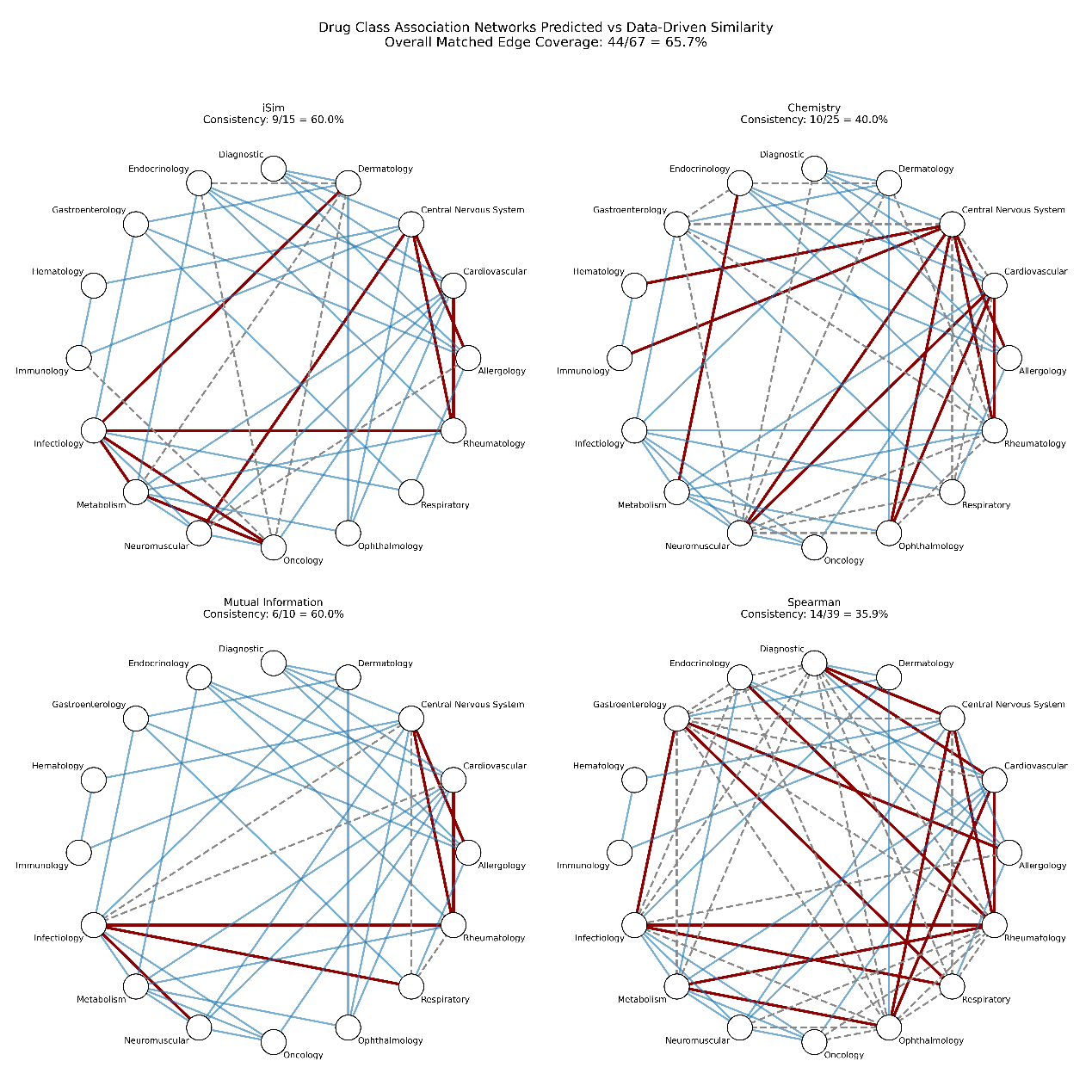

**Consensus similarity network of drug categories.** The network compares TopoMPI-C–predicted similarities with established pharmacological similarity metrics, including iSIM (integrated similarity), chemical structure similarity, mutual information, and spearman correlation. Nodes represent drug categories; edges indicate predicted similarity, with edge colors denoting consistency with external benchmarks.

**Supplementary Figure 9**

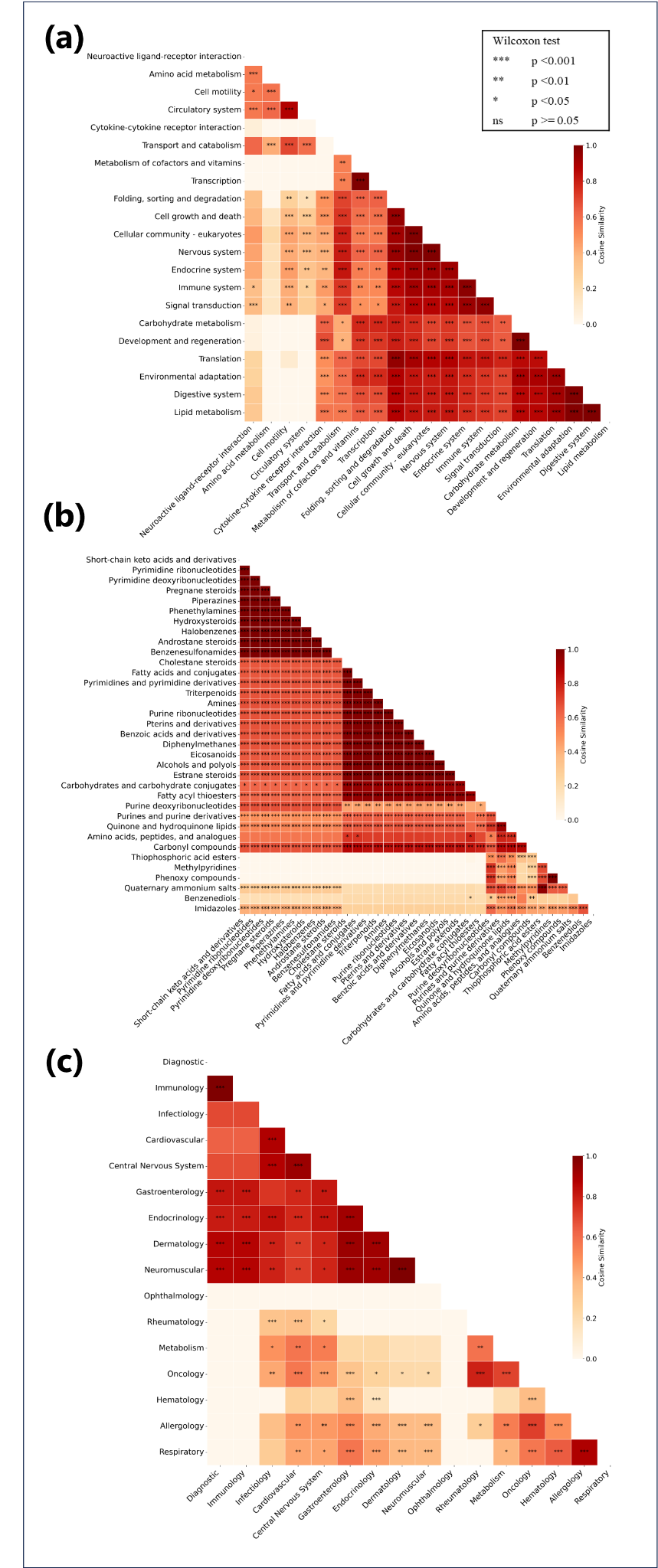

**Cosine similarity heatmaps derived from the Sankey diagram**, representing protein pathways (a) , metabolite classes (b) and drug classes(c). Axes denote category labels; color intensity indicates cosine similarity based on the respective group’s interaction profiles. Drug category similarity shows strong intra-class clustering and functional convergence; pathway similarity reveals consistent reuse across drug categories, suggesting common regulatory targets; and metabolite class similarity exhibits modular patterns, confirming TopoMPI-C’s ability to capture structured, cross-layer biochemical organization.

**Supplementary Figure 10**

**Integrative analysis of TopoMPI predictions for histone-modifying enzymes**. Panels a–d correspond to four major histone-related enzyme categories: acetylases (a), deacetylases (b), demethylases (c), and methylases (d). In each panel, the left subfigure shows unique predictions (edges or triplets supported by only one model), where TopoMPI-D, -I, and -C are distinguished by blue, green, and orange edges, respectively. The right subfigure shows the consensus matrix of metabolite–protein pairs predicted by multiple models, with cells color-coded by the specific combination (DI, DC, IC, or DIC). Nodes represent proteins, metabolites and drugs. All metabolite and drug labels shown in red font correspond to epigenetic metabolites or drugs. Unique predictions capture complementary aspects of direct metabolic inputs, indirect redox- and signaling-mediated links, and drug-conditioned regulation, while consensus predictions highlight robust and recurrent biochemical principles. Collectively, these results demonstrate how TopoMPI integrates complementary perspectives to uncover both established and novel metabolic–epigenetic regulatory axes.
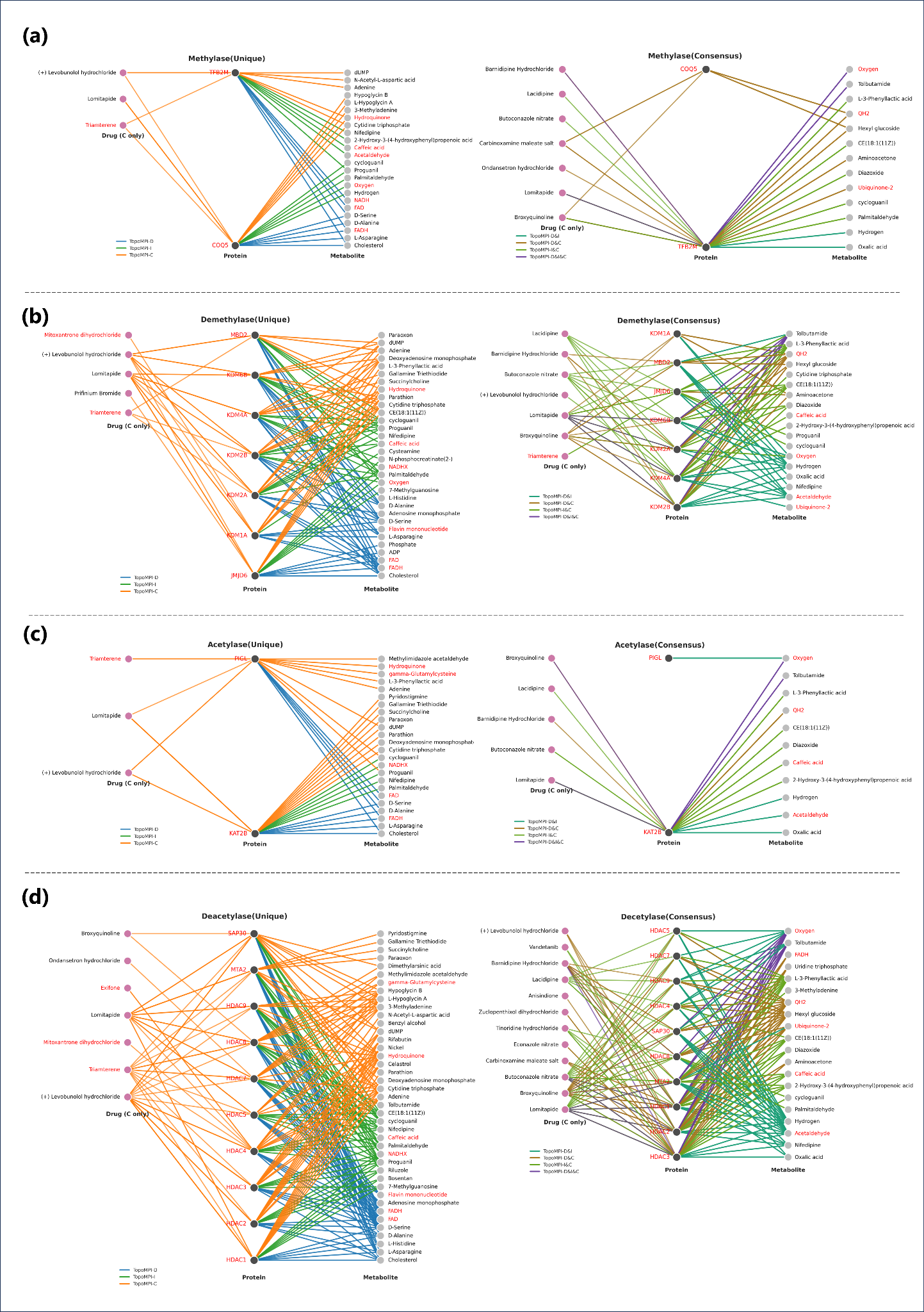

**Supplementary Tables**

**Supplementary Table 1**

Centrality ranks of protein nodes across 24 tissue-specific networks in the original network and three extended networks. *Tissue* represents the tissue-specific network; *node* represents the protein node ID; *Original* is the centrality rank in the original network; *Pred_top_5000*, *Pred_top_10000*, and *Pred_top_15000* are the centrality ranks in the extended networks with the top 5k/10k/15k predicted edges added, respectively. *delta_5000*, *delta_10000*, and *delta_15000* represent the rank changes compared with the original network. The detailed table is provided in Supplementary_Table_1.csv.

**Supplementary Table 2**

Centrality ranks of metabolite nodes across 24 tissue-specific networks in the original network and three extended networks. Column definitions are the same as in Supplementary Table 1, except that *node* represents metabolite IDs.The detailed table is provided in Supplementary_Table_2.csv.

**Supplementary Table 3**

Reachability scores of KEGG pathways across 24 tissue-specific networks in the original network and the extended network. *Tissue* represents the tissue-specific network; *Pathway* represents the KEGG pathway ID; *Original* is the reachability score in the original network; *Pred_top_15000* is the reachability score in the extended network; *Reach_diff* represents the score change between the two networks; *Pathway_info* provides detailed pathway annotations.The detailed table is provided in Supplementary_Table_3.csv.

**Supplementary Table 4**

Tissue–pathway pairs with improved connectivity after TopoMPI-D prediction. *Pathway_info* represents the KEGG pathway information;*Tissue* represents the tissue-specific network; *Avg_reach_diff* represents the score change between the two networks. The detailed table is provided in Supplementary_Table_4.csv.

**Supplementary Table 5**

Predicted metabolite–protein and metabolite–protein–drug associations involving 21 histone proteins jointly identified by all three TopoMPI submodels. The table includes the following columns: *protein* (UniProt identifier), *protein_category* (histone enzyme class: methylase, demethylase, acetylase, or deacetylase), *metabolite* and *metabolite_name* (HMDB or KEGG identifier and corresponding metabolite name), *drug* (drug name, reported only for TopoMPI-C predictions), *model* (TopoMPI submodel that generated the prediction), *pred_score* (prediction score assigned by the model), and *model_count* (number of submodels predicting the same metabolite–protein interaction). The detailed table is provided in Supplementary_Table_5.csv.

**Supplementary Table 6**

Literature-supported and putative metabolite–drug associations derived from the TopoMPI leave-one-drug-out (LODO) analysis. The table summarizes, for the ten representative drugs in the LODO setting, the top 20 prioritized metabolite candidates per drug (200 pairs in total). The following columns are reported: *drug* (drug name), *metabolite_name* (common name of the metabolite), *priority* (normalized composite priority score used in Fig. 6c–d), and three evidence-annotation columns, *direct*, *indirect*, and *potential*, which record the primary type of literature support identified by manual curation. Direct evidence denotes publications explicitly reporting drug-induced changes in the metabolite; indirect evidence captures mechanistic or pathway-level links involving both entities; potential evidence indicates co-mention or contextual association without explicit causal statements. Entries with no annotation in any evidence column are classified as novel predictions. The detailed table is provided in Supplementary_Table_6.csv.
